# Judging the reasons for fixations: A direct experimental method to assess the contribution of saliency and semantic factors to gaze control

**DOI:** 10.64898/2026.07.01.735892

**Authors:** Franz Faul, Antje Nuthmann

## Abstract

Current debates regarding the relative contribution of saliency versus semantics to gaze control often rely on comparing the predictive power of saliency and meaning maps. We argue that such indirect, global approaches are fundamentally limited because fixations arise from heterogeneous, local causes that are conflated in whole-scene comparisons. To substantiate this claim, we used a direct method where participants explicitly identified the reasons for fixation at specific clusters of high fixation density, distinguishing between low-level saliency and various semantic categories, as well as the most important one. The obtained judgments revealed that multiple factors contribute simultaneously to gaze control. Although their influence varied across fixation clusters, semantics generally dominated saliency. Notably, abstract semantic categories, particularly “unknown/unusual,” proved important, highlighting the role of prior knowledge and novelty besides personal relevance in guiding attention. To interpret these findings in the context of existing models, we propose a framework distinguishing between processes highlighting interesting locations in the image from a sampling strategy translating this information into scanpaths. Within this framework, classic saliency and meaning maps are viewed as restricted inputs to the strategy, whereas deep learning-based models (e.g., DeepGaze IIE) are more general and may also implicitly encode aspects of the strategy itself. Consistent with this, we found that the predictive performance of DeepGaze IIE varied less significantly with the specific reasons for fixation than that of classic saliency and meaning map approaches.

## 1 Introduction

When an observer views a natural-scene image without an explicit task or goal, fixations are not randomly distributed. Instead, when fixations are aggregated across many participants, they converge on a spatial pattern that is unique to each image. These fixation distributions are typically sparse, comprising only a few hotspots (Figure 1).

**Figure 1:**
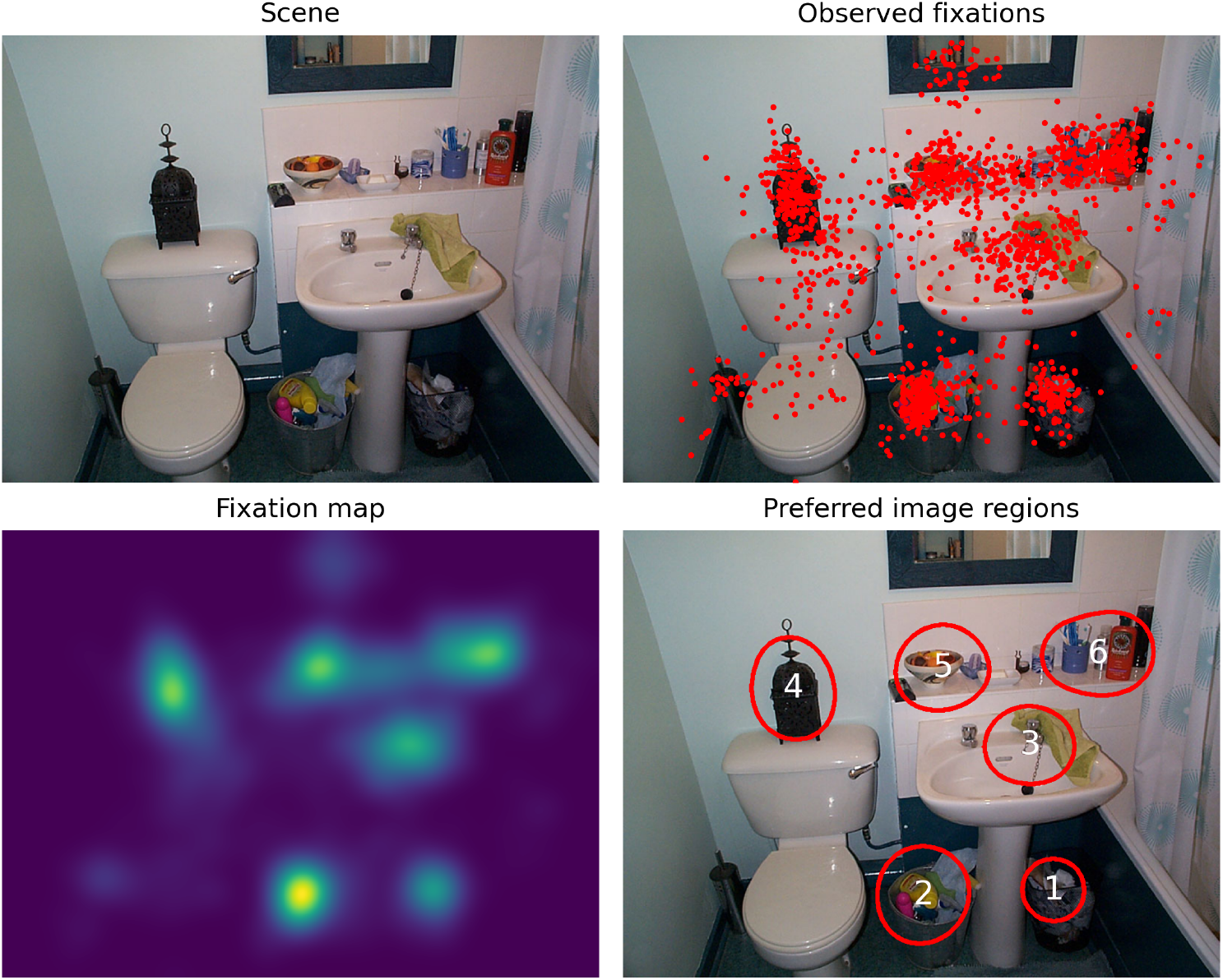
One of the scenes used in the study (top left). The fixations of different observers cluster in a few areas of the image (top right). Smoothing the raw fixation locations with a Gaussian results in a fixation map (bottom left). Setting the values in the fixation map below and above a threshold value to zero and one, respectively, defines a mask for the most relevant regions. The red outlines in the bottom-right panel show the boundaries of expanded versions of such regions.

Two primary theoretical frameworks—referred to as image guidance theory and cognitive guidance theory by Henderson and Hayes (2017)—have been proposed to account for this regularity. Both theories posit that eye fixation patterns reflect the spatial distribution of overt attention across a visual scene. Image guidance theories hold that attention is drawn involuntarily to visually salient features, such as edges, contrasts, or other low-level perceptual cues (Borji & Itti, 2013; Foulsham, 2019; Tatler et al., 2011). In contrast, cognitive guidance theories emphasize the observer’s active role, proposing that fixations are directed toward image regions that are meaningful or task-relevant (Henderson, 2020a; Henderson et al., 2009; Underwood et al., 2006; Walter et al., 2024). For brevity, we henceforth refer to these two influences as saliency (image-driven) and semantics (cognition-driven), respectively.

Research on eye movements during picture viewing has evolved substantially in both aims and terminology over time. Early investigations emphasized higher-order cognitive processes, focusing on the types of objects that observers fixated (Buswell, 1935; Yarbus, 1967). In these early approaches, sequences of recorded fixations were regarded primarily as a means of identifying the objects within a scene that were considered important or meaningful by the observer.

The introduction of the influential feature-integration theory of attention (Treisman & Gelade, 1980) marked a significant shift towards the influence of low-level image features on gaze control. Building on this framework, the seminal contributions of Itti et al. (1998) and Itti and Koch (2000) catalyzed the development of a growing body of computational models aimed at estimating spatial saliency distributions from natural-scene images. These models are typically validated by assessing how accurately the resulting saliency maps predict human fixation distributions (Kümmerer et al., 2018). Over subsequent years, particularly within the computer vision literature, research progressively shifted from identification of attended objects to the prediction of fixation locations and even complete scanpaths using machine learning (Kümmerer & Bethge, 2023). This development contributed to a gradual conflation of the concepts of “semantics” and “saliency”. In much of the recent literature, saliency has come to be “directly equated with fixation probability” (Kümmerer & Bethge, 2023, p. 275). Conversely, Henderson et al. (2019) introduced new conceptual distinctions by using the terms “semantic salience” in contrast to “image saliency” and by defining a “semantic feature” (p. 1).

The two explanatory frameworks are not mutually exclusive. Under natural viewing conditions—or when naturalistic stimuli are used—it is plausible that both bottom-up processes driven by image saliency and top-down processes guided by scene semantics jointly influence gaze behavior. However, the relative strength of these processes likely depends on factors such as viewing duration, scene content, and the observer’s prior experience. This raises the question of how to quantify the respective influence of these factors in a given context. In the present work, we propose and evaluate a direct experimental method to address this question. To motivate this approach, we first review previous methods and discuss their potential limitations.

### Existing approaches for evaluating the contribution of saliency and semantics

A well-established strategy for evaluating the role of saliency in gaze control involves computing a saliency map—the spatial distribution of saliency across the scene—using a chosen saliency model. If saliency influences fixation behavior, fixation densities should be higher at locations with greater saliency values. Consequently, a strong correspondence between saliency and fixation maps indicates that the information encoded in the saliency map substantially influences gaze control. Various metrics have been proposed to quantify the similarity between the spatial distributions of saliency and observed fixations (Bylinskii et al., 2019; Kümmerer & Bethge, 2023). Several known issues with this approach—most notably, the dependence of the results on the selected metric—were addressed by the systematic evaluation framework introduced by Kümmerer et al. (2018), which proposes transformations that optimize the performance of a map for a specific metric. An alternative approach uses generalized linear mixed models to compare the performance of saliency models in predicting fixation selection (Nuthmann et al., 2017).

To enable a comparable validation strategy with respect to cognitive guidance, Henderson and Hayes (2017) introduced meaning maps as a semantic analog of saliency maps (see also Henderson et al., 2019). A meaning map for an image is generated by extracting partially overlapping patches arranged on a regular grid at two spatial scales (“fine” and “coarse”). Participants then rate each patch for the extent to which it contains meaningful information; these ratings are averaged and spatially smoothed, producing a continuous distribution of “meaning”. To demonstrate how meaning maps can be used to assess the relative contributions of semantic and saliency information, Henderson and Hayes (2017) correlated observed fixations for 40 images with (a) meaning maps and (b) saliency maps generated using the GraphBased Visual Saliency (GBVS) model (Harel et al., 2007). The correlations were significantly higher for meaning maps, leading the authors to conclude that semantic information plays a greater role than saliency in gaze control. A similar pattern was reported using “contextualized” meaning maps, in which raters saw not only the image patch but also its location within the image during the rating procedure (Peacock et al., 2019).

However, conclusions about the relative importance of saliency and semantic information drawn from such map comparisons may be misleading if one or both maps fail to capture the full range of the respective information relevant for gaze control (Leemans et al., 2024; Pedziwiatr et al., 2021a). Determining whether this is the case is often difficult, as both saliency and semantic information are represented only implicitly and in a combined form within the respective maps. This problem also affects the approach by Kümmerer et al. (2017) to compare models with low- and high-level feature spaces to investigate the relative contribution of low- and high-level information.

Classic saliency models, such as GBVS, generate saliency maps algorithmically from the scene image. Thus, contributing factors are explicitly defined, and their relative influence can, in principle, be assessed by selectively adding or removing individual components. By contrast, saliency models that rely on machine-learning techniques, such as DeepGaze IIE (Linardos et al., 2021), are trained on empirical fixation data; consequently, the relevant factors are not explicitly known, and the models are harder to interpret (Borji, 2021). Moreover, it is likely that both low-level saliency and high-level semantic aspects contribute jointly to the resulting saliency map.

Meaning maps are particularly problematic in this respect, as their very construction means that they capture only a small fraction of the semantic information known to influence gaze control. In the rating procedure, participants are instructed to “assess the meaningfulness of each patch based on how informative or recognizable they thought it was” (Henderson & Hayes, 2017, p. 4). Consequently, ratings of isolated image patches primarily reflect the extent to which their content can be recognized without reference to the broader scene. This holds even with contextualized meaning maps (Peacock et al., 2019), because the additional information provided in this case is intended only to aid raters in judging whether the local information available inside the patch suffices to recognize the actual scene content.

This approach introduces several methodological difficulties. Because the patch size is fixed, relatively homogeneous objects that extend over several patches may receive low ratings, as most patches covering the object appear uninformative. Conversely, it is often unclear how to rate a patch containing easily recognizable structures, such as grass in a meadow, that are presumably of little relevance to an observer. More fundamentally, meaning maps cannot capture semantic information that is task-dependent or relational in nature, such as an object being incongruent with the overall scene context (Henderson et al., 2021; Pedziwiatr et al., 2021a, 2022). Like many saliency maps, they also fail to capture that vague or ambiguous information can itself be a source of interest and attract attention.

Most of these limitations arise from the indirect nature of these methods, which infer the relevance of a potential factor—be it saliency or meaning—solely from its ability to predict observed data. Indirect measures of psychological phenomena are often regarded as more objective than direct reports; however, this advantage only holds when the relationship between the indirect measure and the underlying phenomenon is well-understood and stable. The diversity of existing saliency models and the evident limitations of meaning maps suggest that this condition is not met. As a result, it remains unclear to what extent these maps capture all perceptually relevant aspects of saliency and semantics, or how much they conflate the two.

### A direct method to determine the reasons for fixations

In this article, we explore a direct experimental method based on the assumption that if we know *where* people prefer to look, we can often infer plausible reasons for their specific interest. This ability was likely what enabled Yarbus (1967) to correctly infer many general principles of semantic interpretation, simply by inspecting the fixation data his participants produced while viewing images. For example, he concluded that the eyes of a person are of special interest and that fixations depend on the given task. It was later shown that task information can be decoded from eye movements in an “inverse Yarbus process” (Boisvert & Bruce, 2016; DeAngelus & Pelz, 2009; Haji-Abolhassani & Clark, 2014).

The application of the proposed direct method requires that fixation data for a set of natural scenes be available in advance. The core idea is to use this data to identify regions of particular interest in the images. These regions are then presented in an experiment in which participants first rate how well each region fits into a set of predefined categories related to saliency and semantics. In a second step, they indicate which of these categories they consider most important for explaining the fixations.

To preview the results, the ratings and categorizations obtained in the experiment described above were highly reliable and appear well suited for assessing the relative importance of the different categories in gaze control. The data indicate that both saliency and semantics encompass several aspects that need to be differentiated. Moreover, several of these factors contributed to each fixation, but the relative weight of each factor varied both within a scene and across scenes. Overall, the results support previous findings suggesting that semantic information plays a greater role than saliency in guiding gaze.

In the second part of the article, we examine the relationship between this direct method and existing indirect methods. To this end, we outline a framework describing the flow of information during gaze control. This framework is then used to identify the specific information captured by different approaches that aim to explain the fixation behavior in participant-directed scene viewing tasks, such as free viewing or scene memorization (Mills et al., 2011). Ultimately, we argue that the direct method yields unique insights not readily accessible from saliency or meaning maps.

## 2 Experiment

The indirect approaches related to saliency and semantics mentioned above condense the relevant information into a single dimension, representing local saliency or local meaning, respectively. In the case of saliency, this seems unproblematic, because different sources of saliency are likely additive. Indeed, this linearity assumption is often explicitly made in saliency models (e.g. Harel et al., 2007) when the contribution of different feature maps are combined into a single map, as well as when generalized linear mixed-effects models are used to identify image factors contributing to fixation selection (Nuthmann & Einhäuser, 2015). The situation is more complicated for semantic information, as semantic attributes can be of very different kinds and may even be partially inconsistent and contradictory (Spotorno & Tatler, 2017). Thus, condensing semantic information into a single dimension appears inadequate.

We argue that it is necessary to distinguish between different aspects of saliency and semantics that (a) can occur independently, (b) are conceptually distinct, or (c) are even mutually exclusive. Adopting this more complex framework implies that determining the relative contributions of saliency and semantics is only one component of the broader challenge of identifying the specific combination of factors directing gaze to a particular location.

The proposed direct method for assessing the contributions of saliency and semantics therefore not only requires to identify the most relevant image regions, but also to provide a concise description of potential causes of fixations. Identifying all relevant factors that contribute to gaze control and organizing them into a limited set of meaningful categories is a challenging and as yet unsolved task. The categories used in our study provide a pragmatic solution that can be refined in future work. We distinguish two aspects of saliency: “visual saliency” and “favorable image position”. Visual saliency refers to low-level image properties—such as contrasts in color, hue, or texture—that cause a region to stand out from its surroundings. In contrast, a favorable image position is one that is, a priori, more likely to be fixated. The most prominent example is the central fixation bias, that is, the tendency to fixate locations near the scene center more frequently than locations farther from the center (Tatler, 2007). We consider this factor part of saliency, since it reflects a non-semantic influence on gaze.

We consider semantic information as either *object-related, task-related*, or *abstract. Object-related semantics* concern important object categories, which can be either innate and universal (e.g., human or animal faces) or culturally learned (e.g., displays, clocks, or text), which are preferentially looked at during scene viewing (Broda & De Haas, 2022; Cerf et al., 2009; Li & Chen, 2021; Wang & Pomplun, 2012). *Task-related semantics* derive their importance from a specific task; objects or environmental states that may aid or obstruct task completion become particularly relevant. While task-related information is crucial in everyday situations, we disregard it in our study, using data from an eye-tracking experiment in which participants performed a memorization task without explicit search or object selection instructions. Finally, *abstract semantic information* involves the relationships between the perceived input and the observer’s prior knowledge. This includes recognizing an object in an unusual location, encountering an unfamiliar object, or finding an image region difficult to interpret due to shadows or partial occlusion. Chakraborty et al. (2022) discuss different concepts of uncertainty and show that object recognition uncertainty has a special role in predicting attention.

In our study, we sought to capture these aspects using nine categories: two for saliency and seven for semantics. The saliency categories were ‘visual saliency’ (optische Auffälligkeit) and ‘favorable image position’ (günstige Bildposition). The semantic categories included four object-related categories—’animal’ (Lebewesen), ‘text or information source’ (Text oder Informationsquelle), ‘food’ (Lebensmittel), and ‘utility object’ (Gebrauchsgegenstand)—and three abstract categories—’unknown or unusual object’ (unbekanntes oder ungewöhnliches Objekt), ‘prominent scene position’ (auffälliger Ort in der Szene), and ‘poorly visible’ (schwer erkennbar). The terms in parentheses indicate the corresponding German category name used in the experiment.

Our set of object-related categories is specifically designed for our stimulus material, which consisted solely of interior scenes. If the method is applied to other scenes, it may be useful to include additional object-related categories, for example, ‘vehicles’ for outdoor scenes.

### 2.1 Methods

#### Stimuli

Scene images and corresponding fixation data were taken from an earlier study (Nuthmann et al., 2020). In the experiment, participants performed a memorization task, with memory probed in a subset of trials by questions about the presence or absence of specific objects in the scene. The fixation data from 42 young adults (mean age 22.1 years) and 34 older adults (mean age 72.1 years) for 150 color photographs of natural scenes were aggregated across age groups. In this eye-tracking study, the images were displayed on a 21-inch CRT monitor with a resolution of 800× 600 pixels that subtended 25.78°× 19.24°.

After excluding the initial central fixation in each trial, raw fixation maps were computed for each scene *S* and then convolved with a Gaussian kernel (σ = 20 pixels, corresponding to roughly 0.64°) to obtain smoothed fixation maps *F M*_*S*_ (see Figure 1). All maps were scaled to a maximum of 1. Masks *M*_*S*_ for preferred image regions (“hotspots”) were calculated by setting pixels *M*_*S*_ (*x, y*) to 0 if *F M*_*S*_ (*x, y*) *<* 0.48 and to 1 otherwise. The threshold value 0.48 was optimized interactively, with the goal to capture only prominent hotspots in the chosen subset of images. To account for inaccuracy in the fixation data, a mask variant *ME*_*S*_ was used, in which the region sizes were expanded by applying the `dilation` function in the `skimage.morphology` module of the Python package `scikit-image`, using a disk-shaped structuring element with a radius of 20 pixels, to *M*_*S*_ (*x, y*) (see Figure 1). We then selected 50 indoor scenes according to the following criteria: (1) the scenes contain at least two preferred regions, (2) the scenes are moderately cluttered, (3) the room types ‘living room’, ‘kitchen’, ‘bathroom’, and ‘garage’ are present in approximately equal proportions, and (4) all semantic categories are covered. From each of the 50 images, two regions with no overlap with (expanded) neighboring regions were selected. A further selection criterion was again that, overall, the regions should cover the full range of our preselected categories.

For each target region, a square sub-image of 200×200 pixels was extracted from the original image, which in each case contained the entire target region. Its position was chosen such that its center minimized the distance to the center of gravity of the target region. The size of these sub-images corresponds to about 6.5°× 6.5° in the original eye-tracking study. In the current experiment, these sub-images were increased in size by a factor of two and shown beside the original image (see Figure 2). The scenes were prepared in two variants. In the ‘without context’ condition, only the expanded region to be assessed was marked with a red outline, whereas in the ‘with context’ condition, the remaining preferred regions were additionally marked with a yellow outline. The rationale for this design was that knowing which other regions were fixated in the previous study could influence which potential reason for fixation was considered the most relevant. For instance, an attribute in the target region is presumably less likely to be considered the most relevant if it is known to be shared with non-fixated parts of the image.

**Figure 2:**
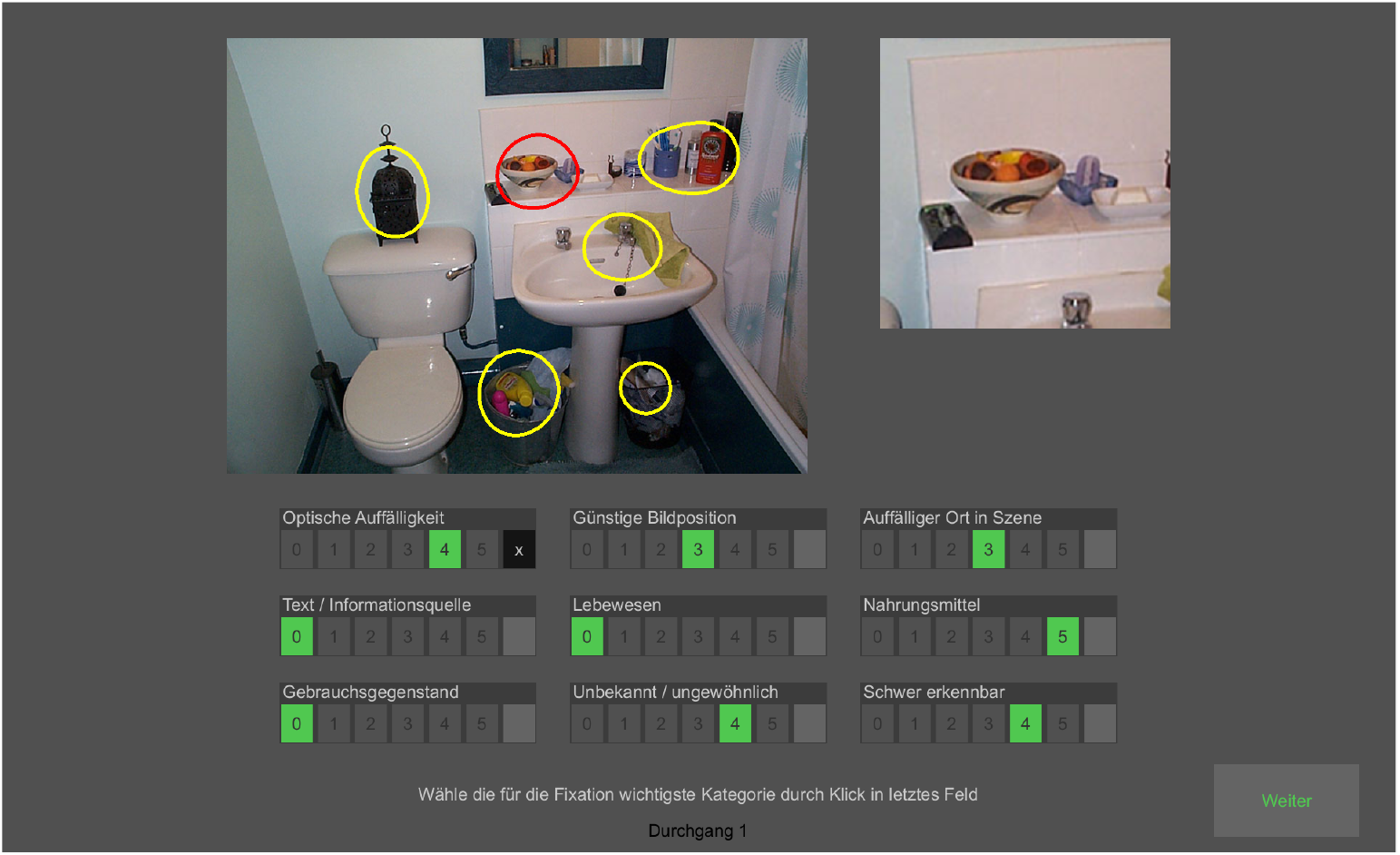
Screenshot of a display during the experiment. The left image (800×600 pixels) shows the scene with marked regions of interest. The red outline marks the region the participants had to judge. The yellow outlines were only visible in the ‘with context’ condition. The smaller image (400 × 400 pixels) on the right shows an enlarged view of a 200 × 200 pixel section of the scene, centered at the centroid of the region in question. Below these images, the scales for each category are shown. The screenshot is from the second phase of the trial, in which the ratings for each category made in the first phase are marked in green. Here, the subject chose the first category as the most relevant one (black cross at the end of the scale).

In the full version of the experiment, a participant performed two blocks. In the first block, one of the regions per image was rated in one context condition. In the second block, both the region and the context condition were different. In this way, all 100 regions were assessed, with each of the two regions per image evaluated under a different context condition. Combining the data of two participants covers all four combinations of region and context, while a full balancing of the order of both conditions can be achieved by combining the data of eight participants. We planned for three fully balanced condition combinations, comprising 24 sessions (48 blocks of 50 regions).

Performing the full version took a considerable amount of time. To facilitate participation, we also offered a short version of the experiment, which consisted of only one block. In this case, two participants together contributed the equivalent of one ‘full version’.

#### Participants

Of the 38 participants (19–36 years, M = 22.3 years), 13 performed the full version of the experiment and 25 the short version. In total, 51 blocks were conducted. Of these, 3 were incomplete (due to technical errors) and were therefore excluded. As a consequence, the final data were only approximately balanced, with 10 to 13 blocks in the four combinations of region and context conditions, and 25 and 23 blocks for the two regions.

#### Procedure

The stimuli were displayed on a 24-inch LCD monitor (Eizo ColorEdge CG243W) with a resolution of 1,980× 1,200 pixels. To optimize perceived contrast and avoid reflections, the experiment was conducted in a darkened room. We used Processing 4.0 (https://processing.org) to implement the experiment.

Figure 2 shows the general outline of the different elements seen by the participants in each trial. In the first phase of each trial, participants assessed the relevance of each of the nine categories for the target region on a scale from 0 to 5, where 0 indicated ‘irrelevant’ and 5 indicated ‘highly relevant’. All choices were made with a mouse click. Clicking on the corresponding screen location gave visual feedback of the selection made. No time restrictions were applied, and participants could change their ratings at any time. When all nine categories were rated, a ‘Continue’ button appeared, which, when pressed, led to phase 2. In this second phase, participants were asked to select the category they considered the most important reason for fixating that region. After a category was chosen, a ‘Continue’ button appeared. Again, the selection could be changed until this button was clicked.

In the instructions immediately prior to the experiment, participants were informed about the exact meaning of each category and their tasks in each trial. The instructions were also available in written form during the experiment. For the object-related categories, participants were asked to base their judgment on how prototypical the depicted object(s) are for the corresponding category.

In total, 2,400 trials were conducted, 1,250 for region 1 and 1,150 for region 2.

### 2.2 Results of the rating experiment

The analysis of the rating data comprises seven parts addressing the following questions: (1) How reliable were the ratings of category relevance and the selection of the most important category? (2) How many categories were considered relevant for a single region? (3) How strongly do the ratings of different categories correlate with one another? (4) How often was each category considered relevant? (5) What is the relative importance of the nine categories? (6) Are saliency or semantic categories overall more important? (7) How can the relationship between category ratings and the selection of the most important category be modeled?

The results for each of these questions are presented together with a brief discussion. A broader overview, including the relation to existing indirect approaches, is provided in the General Discussion.

#### Inter-rater reliability

A central assumption of the direct approach is that it is possible to infer the reasons why people prefer to look at a certain location. A minimal requirement for this assumption to be valid is that different observers generally agree on the relevant aspects of the image content at this location. If this holds and the nine categories adequately cover the relevant aspects of image content, one would expect a considerable degree of agreement in the ratings. For ordinal rating scales, intraclass correlation (ICC) is a suitable measure of agreement between raters. We computed ICC using the R package *icc* (Gamer et al., 2019), specifying a two-way model and single-measure ICC across the 50 scenes. Figure 3A shows the consistency ICC for each of the nine categories, separately for the two regions. Unlike the absolute-agreement ICC, the consistency ICC ignores differences in overall rating level and is therefore more appropriate in this context. Nevertheless, for the present data, the absolute-agreement ICC was only slightly lower and exhibited the same pattern. Kendall’s coefficient of concordance *W*, another alternative for ordinal data, produced almost identical results (not shown).

**Figure 3:**
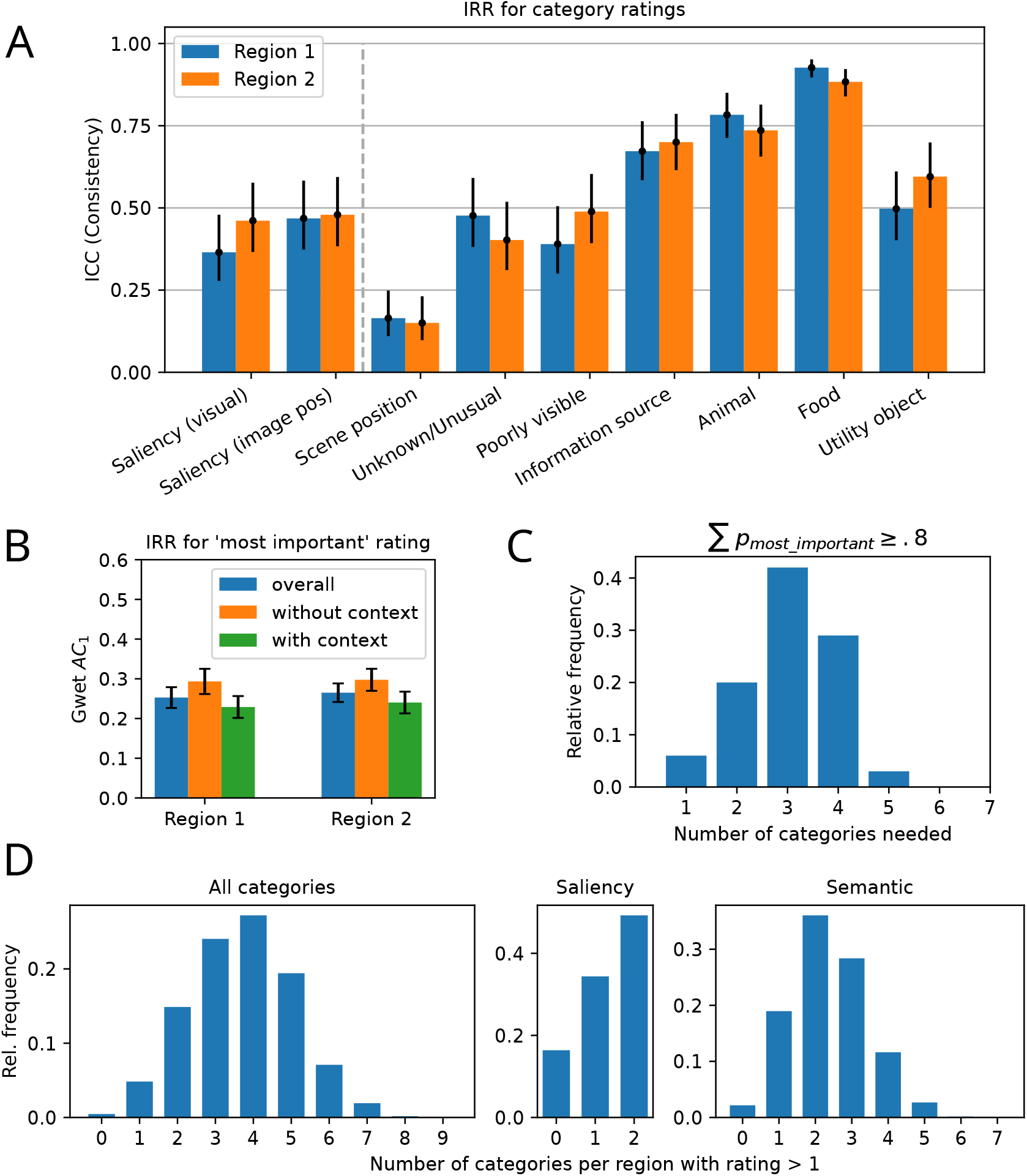
Inter-rater reliability (IRR) and number of categories used per region. A) Intraclass correlation for each category, computed separately for region 1 (*n* = 25 raters), and region 2 (*n* = 23 raters). Error bars indicate the 95% CI. B) Inter-rater reliability in judging the most important category. Gwet’s *AC*_1_ is shown for the two regions; within each region, values are shown for the two context conditions and overall. C) Distribution of the minimal number of categories with a cumulative probability of being the most important category ≥ .8. D) Relative frequency of the number of categories per region with a rating greater than 1. Overall, 8,877 of the 21,600 ratings (41.1%) had a value > 1. Left: for all nine categories; middle: for the two saliency categories; right: for the seven semantic categories.

**Figure 4:**
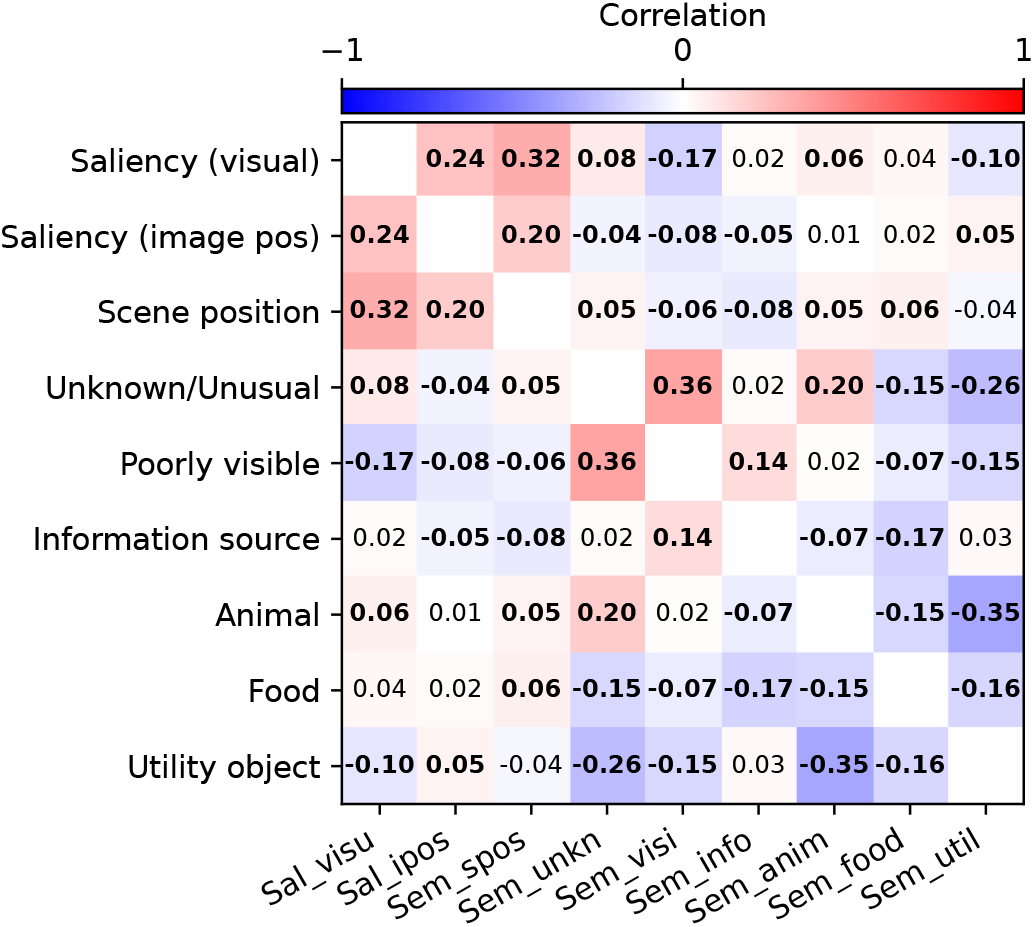
Pairwise correlations between category ratings. Red and blue indicate positive and negative correlations, respectively, and bold numbers indicate significant deviations from zero (*α* = 0.05). *Note*. The *x*-axis labels are abbreviated versions of the corresponding *y*-axis labels.

According to Koo and Li (2016), the observed ICC values correspond to moderate to good reliability. The only exception is ‘scene position’, for which low reliability was found. The highly similar profiles across the categories for both regions indicate true differences between the categories. An interesting observation is that the reliability in judging saliency, which seems to be a relatively simple task, is not higher than that for the semantic categories.

A similar requirement for inter-observer agreement can also be imposed for the second task, namely selecting the category that appears most important for the fixation. For this nominal scale, we used Gwet’s *AC*_1_ (Gwet, 2014) to measure the correspondence between multiple raters. The calculation was done with the Python package irrACA. Figure 3B shows the corresponding values for each region and context condition. Again, the results for both regions are similar. The context was intended to provide additional information about the observers’ general interest; however, surprisingly, the agreement tends to be slightly *larger* without context than with context. A possible explanation is that only some participants used this additional information. Nevertheless, as the effect of context is small in absolute terms, we pooled the data from the two context conditions for further analyses.

According to the benchmark proposed in Gwet (2014), the obtained values between 0.2 and 0.3 correspond to a fair strength of agreement on the Landis-Koch scale. However, considering (a) the relatively large number of categories, (b) the lack of rater training, and (c) the possibility that several categories may contribute to a similar degree, the correspondence seems quite substantial. The last point is particularly relevant, because *AC*_1_ and similar measures like Fleiss’ κ implicitly assume a single correct category, meaning any variation in ratings is attributed solely to rater error. Although our experimental design forced each participant to choose a single category as the most important one, the relevance ratings suggest that several categories contribute to the fixation with similar strength. In this case it is expected that choices will vary between raters. Figure 3C demonstrates that this is indeed the case. It shows the relative frequency of category combinations of size *n*, whose total probability of being the most important category is ≥ .8. If at least 80% of the participants agreed on a single category, this “consensus depth” *n* would be 1; it would be 2 if at least 80% chose one of two fixed categories, and so forth. The peak of the distribution is at *n* = 3 and its mean 3.03, that is, on average about three categories are needed to fulfill the criterion *p* ≥ .8. We can therefore conclude that often several categories contribute with comparable strength to the fixation. On the other hand, this observed *n* is much lower than the expected value *n* ≈ 7 in a null model, where the subjects just sample possible reasons according to their overall popularity. To compute the expected value under this null model, we sorted the marginal probabilities for each reason shown in the middle panel of Figure 5 in descending order. The cumulative sum, which represents the most concentrated sample under this null model, crosses the 80% threshold at *n* = 7. Comparing the observed values of *n* = 3 with the null baseline of *n* = 7 demonstrates that raters are concentrating their choices far beyond what would be expected by arbitrary sampling.

**Figure 5:**
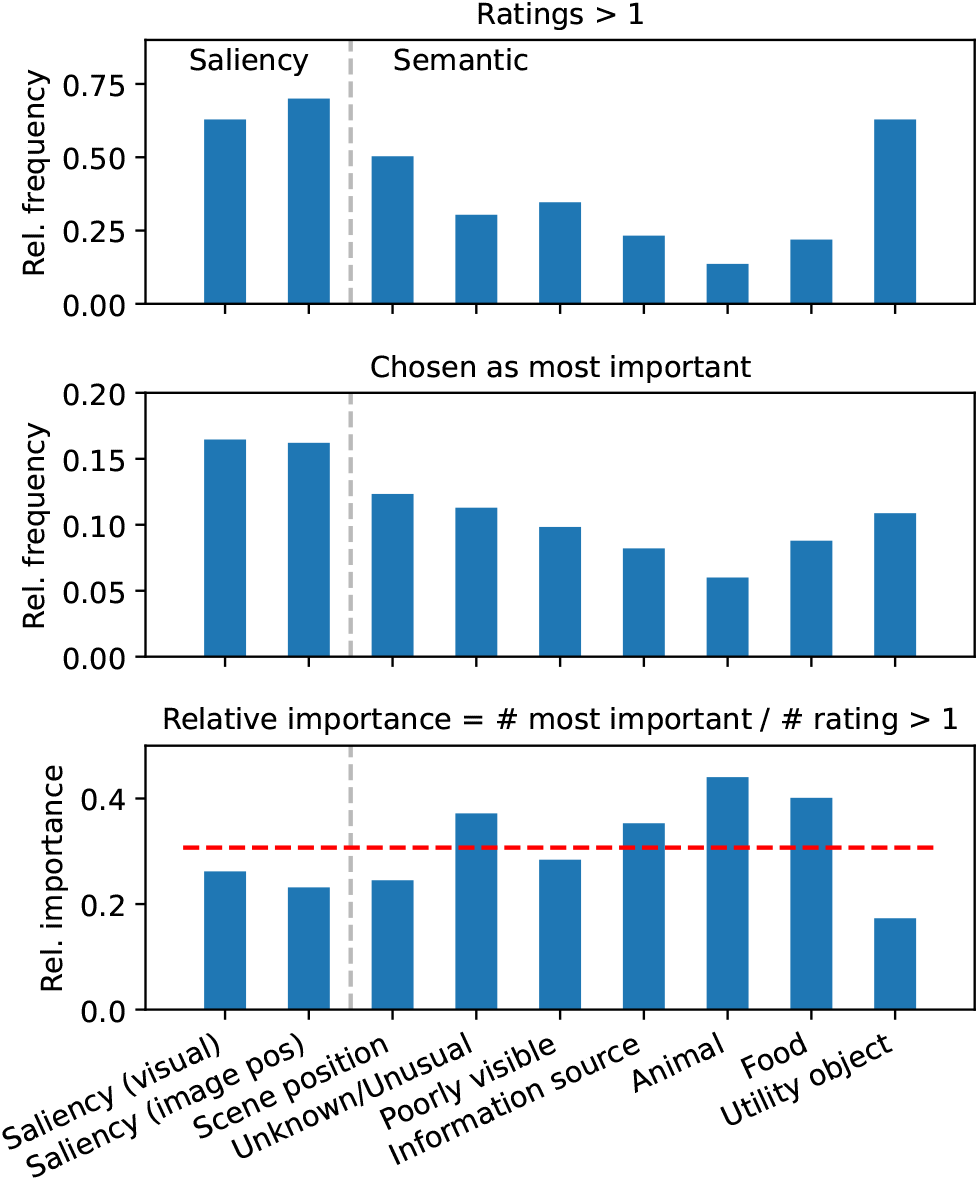
Frequency and importance of the nine categories. Top: Relative frequency of a rating > 1. Middle: Relative frequency of being chosen as the most important category for the fixation. Bottom: Relative importance of each category, defined as the ratio of its frequency of being chosen as most important to its occurrence, i.e., number of ratings > 1. The red line indicates the mean relative importance across all categories.

#### Number of relevant categories

In addition to the assumption that several types of saliency and semantic categories exist, we also assume that several of them are simultaneously active and contribute to fixations at a specific location. We use a rating of 2 or larger as an indicator that a certain category is active. This choice is motivated by the results presented in Figure 9, which show that categories selected as the most important category typically had a rating of at least 2.

Figure 3D shows the distribution of the number of active categories in a single region. If all nine categories are considered together, the peak is at four categories. Regarding the two saliency categories alone, we find that in about 34% of all cases only one of the two saliency criteria was active, and in 16% of the preferred regions neither was active. The “0” cases suggest that saliency is not necessary for a fixation; the “1” cases suggest that the two saliency criteria are independent; and the “2” cases suggest that they can co-occur. For semantic categories alone, the peak is at 2 simultaneously active categories. Interestingly, in less than 0.5% of the cases, no semantic category was active. This suggests that regions that do not convey semantic information are almost never fixated. One reason for the single-peaked distribution is that semantic categories, especially the object-related ones, tend to be mutually exclusive. For example, if an object is classified as food, it is unlikely that it would also be classified as an information source. However, this does not exclude the possibility that it may simultaneously be classified as unusual or poorly visible.

#### Correlation between categories

The relationships between categories are assessed using pairwise correlations between their ratings. Figure 4 shows the resulting Spearman rank correlation coefficients. Two clear trends are visible: the two saliency criteria correlate positively, whereas all object-related semantic categories are negatively correlated, in line with the assumption that they tend to be mutually exclusive. For the abstract semantic categories, a more complex pattern emerges, but there are plausible interpretations for most of the larger correlations. For example, it is plausible that poorly visible objects are often classified as unknown (*r* = 0.35) and that, in such cases, visual saliency tends to be low (*r* = −0.17). The positive correlation between the categories ‘animal’ and ‘unknown/unusual’ (*r* = 0.2) is most likely specific to our scenes, in which animals were often present in unusual forms (e.g., as sculptures, images, or stuffed specimens), and were depicted at large distances or from unusual viewing angles. Similarly, the positive correlation between ‘text or information source’ and ‘poorly visible’ (*r* = 0.14) may at least partially be attributed to the fact that text was often barely legible. A positive correlation between ‘prominent scene position’ and the two saliency criteria (*r* = 0.32 and *r* = 0.20) is also plausible, because this category can be viewed as a hybrid between semantics and saliency. It has a semantic component because it requires at least a partial recognition of the scene; nevertheless, like saliency, selection of this location is largely independent of the semantic properties of the object at that location.

#### Frequency and importance of each category

Figure 5 shows for each category (a) the frequency of receiving a rating *>* 1, (b) the frequency of being selected as the most important category for the fixation, and (c) its relative importance, defined as the ratio of (b) to (a). The observed frequencies of ratings *>* 1 indicate that fixated regions, which are exclusively considered in our study, often appear salient in some form. With regard to semantic categories, this frequency is not particularly informative in itself, as it naturally depends on the type of scenes selected for the sample. This is especially true for object-related categories. The high frequency for the “utility object” category likely stems from it being broader than the other object-related categories. In the present context, the frequency of ratings *>* 1 serves primarily as a reference point to compute the relative importance of each category. Notably, four of the six semantic categories exhibit a relative importance greater than the mean, whereas the values for both saliency categories lie well below the mean. This is a first indicator that semantic factors are more important than saliency.

**Figure 6:**
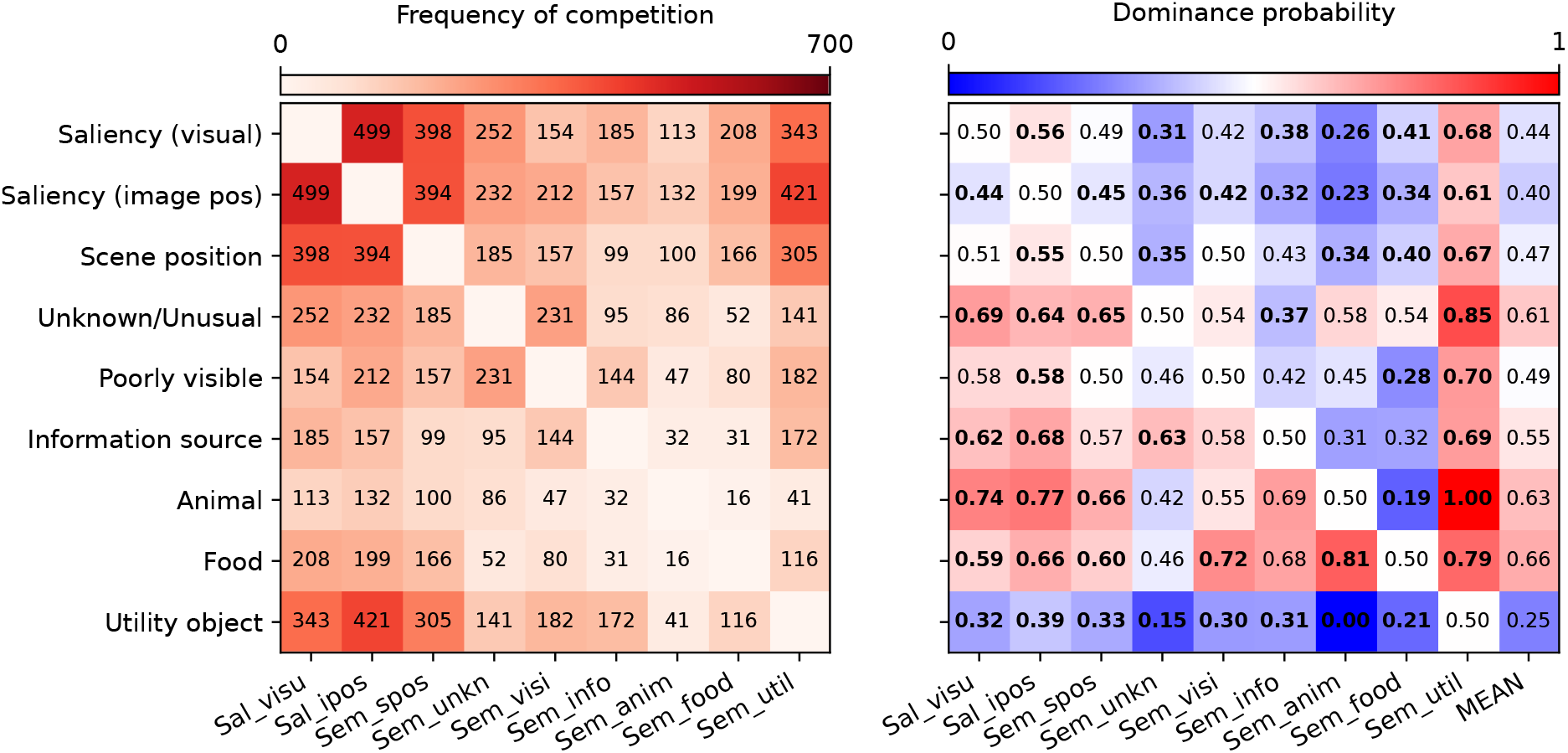
Pairwise dominance of different categories in predicting fixations. Left: Frequency of competition, i.e., the number of category pairs where both received a rating *>* 1 and one of them was chosen as the most important category for the fixation. Right: Estimates of the dominance probability for each category pair. The rows show the dominance of a specific category relative to all other categories. A value *>* 0.5 (red) indicates a dominance of the row category over the column category. The last column gives the mean dominance over the other categories (diagonal elements are excluded from the mean). *Note*. The *x*-axis labels are abbreviated versions of the corresponding *y*-axis labels.

#### Pairwise dominance of categories

A central hypothesis is that there are different types of salience and semantics that may all differ in importance. The results presented in Figure 5 support this hypothesis, but there is an alternative, more direct way to test this. The idea is to estimate the outcome of a competition for fixation between different categories when they are both present in a scene. To this end, we identified for each pair of categories cases in which these two criteria competed. These critical cases were defined by two criteria: (1) both categories *A* and *B* were active (ratings *>* 1), and (2) one of them was chosen as the most important category. We then computed *d* (*A*) = *f* (*A*) /*n* as an estimator of the dominance probability of *A* over *B*. Here *f* (*A*) is the number of critical cases in which *A* was chosen as the most important category and *n* the total number of critical cases concerning *A* and *B*. By construction, *d* (*B*) = 1 − *d* (*A*) and *d* (*A*) = 0.5 means equal importance or no dominance.

The left panel in Figure 6 shows the distribution of the number of critical cases, which is highly uneven across category pairs. This is expected as some categories, for example the two saliency categories, are compatible with some or all other categories, whereas others, in particular the object-related semantic categories, tend to be mutually exclusive. In the latter case, one may even ask why there are any critical cases for seemingly incompatible categories like ‘animal’ and ‘information source’. We address this question below. As the number of critical cases depends on the compatibility of the categories, the frequency matrix is closely related to the correlation matrix: A high positive correlation between categories tends to produce many critical cases, whereas a highly negative correlation results in few such cases.

The observed dominance probabilities are shown in the right panel of Figure 6. Their accuracy is low when based on only a few critical cases, which should be considered in their interpretation. The first observation is that the dominance patterns of the criteria shown in the first three rows are similar. The criterion ‘visual saliency’ and the semantic criterion ‘prominent scene position’ are of similar importance (i.e., *d* (*A*) *~ d* (*B*) *~* 0.5), and both are slightly more important than the saliency criterion ‘image position’. However, all three tend to be less important than the semantic categories shown in rows 4 to 9. The only exception is the category ‘utility object’, which according to this measure is the least important category. This can be easily explained by the observation that ‘utility object’ often appears together with two other object-related semantic categories (see critical cases for ‘utility object’ in the last row in the left panel of Figure 6). A clock would be a typical case of a combination of ‘utility object’ with ‘information source’, while food on a plate represents a combination of ‘utility object’ with ‘food’. The dominance matrix shows that in these cases the more specific category is generally considered more important. Given the small number of critical cases observed for pairs of object-related categories, the corresponding pairwise values are likely of low accuracy. In these cases, the mean dominance shown in the last column may be more informative about their relative importance. Here, both individual and mean dominance values suggest that the (presumably) innate categories ‘food’ and ‘animal’ are the most important object-related categories.

Notably, mean dominance across all categories resembles relative importance (Figure 5). This is illustrated in Figure 7, which compares mean dominance and a scaled version of relative importance.

**Figure 7:**
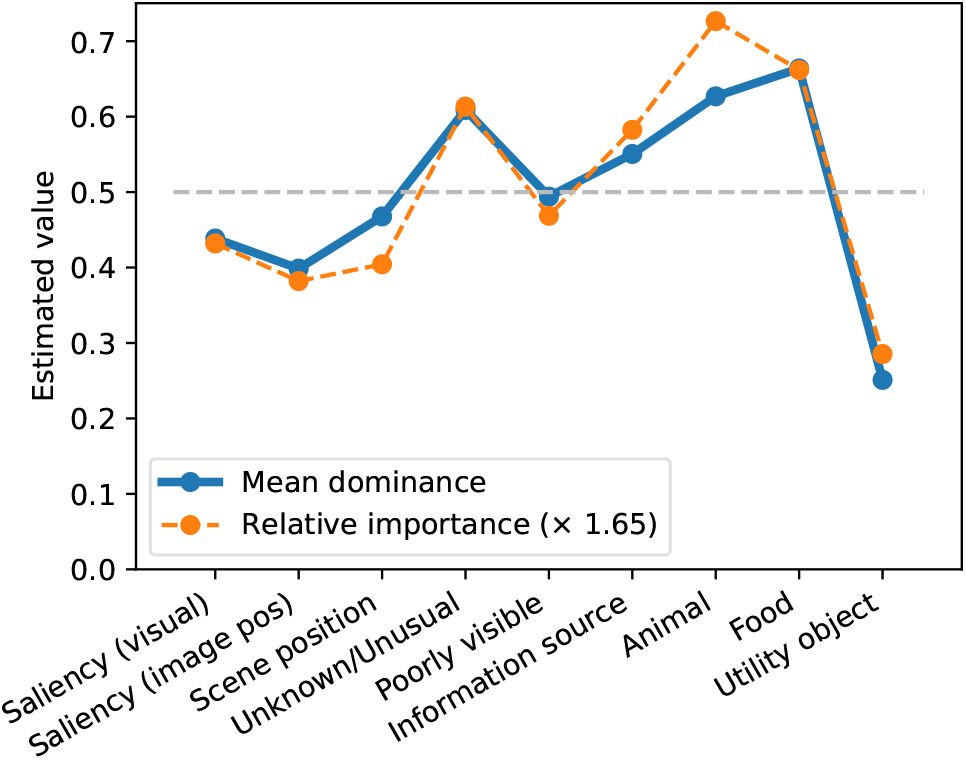
Observed mean dominance vs. relative importance. The pattern of observed mean dominance across categories is similar to the pattern of relative importance (bottom panel in Figure 5).

#### Saliency vs semantics

To determine whether semantics or saliency was overall more important in our stimuli, we collapsed the two saliency categories and the seven semantic categories into saliency and semantics. For each of the 100 regions, we computed the category type *c*_*i*_ with the highest frequency of ‘most important’ ratings, and the category type *c*_*r*_ with the largest mean rating. The categories *c*_*i*_ and *c*_*r*_ are considered consistent if they are of the same type (either both semantics or both saliency); otherwise, they are inconsistent. Figure 8 shows that in 78.1% of the 73 consistent cases, *c*_*i*_ and *c*_*r*_ were both semantic categories. In 74.1% of the 27 inconsistent cases, the saliency category had the highest rating, but a semantic category was nevertheless rated as the most important. Both results support the hypothesis that semantic categories were more important for fixation selection than saliency criteria.

**Figure 8:**
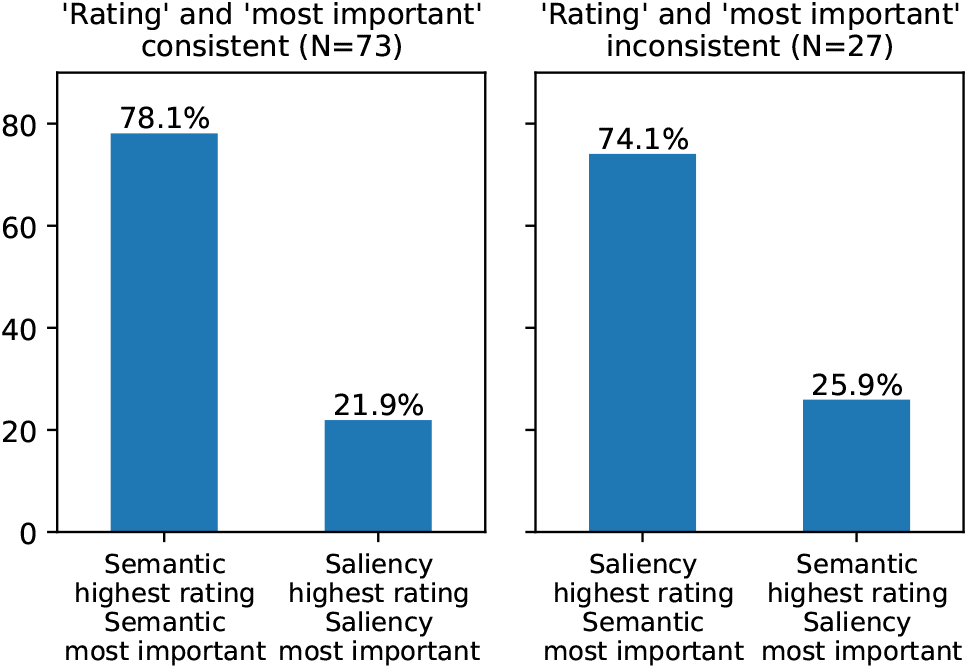
Comparing category ratings and most important judgments for the collapsed categories “saliency” vs. “semantics”. Left: Consistent cases, where the category type with the largest mean rating and the category type with the highest frequency of most important choices is identical. Right: Inconsistent cases.

#### Modeling how the most important category depends on the category ratings

We conclude with an analysis of how relevance ratings for each category are combined to determine the most important category. For each category and each ordinal rating value, Figure 9 shows the probability that the category is chosen as most important. A common trend is that this probability increases monotonically with the rating, which is to be expected if the category genuinely represents a potential reason for fixations.

**Figure 9:**
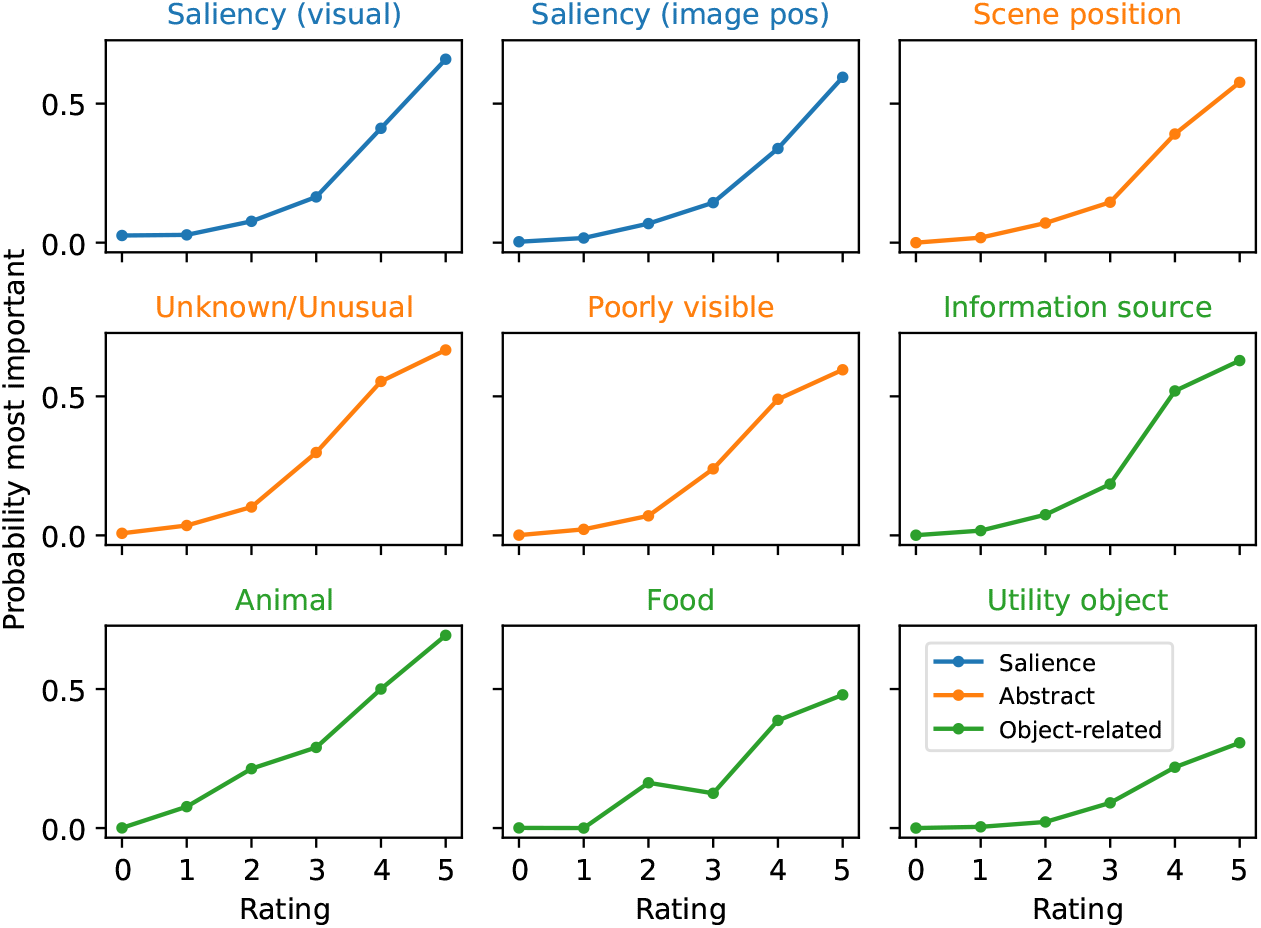
Relationship between category rating and the probability of being chosen as the most important category. For a given category, each data point represents the ratio between the number of cases in which the category was chosen as most important and the total number of cases with the same rating. Thus, a value of 1 for a given rating would mean that all cases with that rating were chosen as most important.

We also fitted a multi-class logistic regression model using category ratings as predictors. This method extends standard binary logistic regression to classification problems with *K >* 2 mutually exclusive outcome categories. In the present case, the task is to select the most important among nine possible categories. The probability of class *k* is given by:

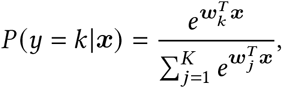

where ***x*** is the predictor vector (the nine category ratings), and ***w***_*k*_ is the weight vector for class *k*. The pre-dicted class *ŷ* is the one with the maximum probability: *ŷ* = arg max_*k*_ *P* (*y* = *k* |***x***).

We implemented the model using the `LogisticRegression` class from the `scikit-learn` library in Python, setting `class_weight = ‘balanced`’. Model performance was evaluated via repeated stratified cross-validation (10 splits, 3 repeats). The mean cross-validated accuracy was 0.62 (*SD* = 0.027), which significantly exceeds chance-level performance (1/9≈0.11).

Figure 10 presents the model results. The left panel shows the confusion matrix, where the entry in row *i* and column *j* indicates the frequency with which observed category *j* was predicted as category *i*. Ideally, all predictions would fall on the diagonal, with off-diagonal elements equal to zero. The observed pattern closely approximates this ideal, confirming good predictive performance. The right panel of Figure 10 displays all weight vectors ***w***_*k*_ from the best fit. The resulting pattern indicates that the probability of selecting a given category as most important depends primarily on the rating assigned to that specific category. The only notable exception is the category ‘utility object’.

**Figure 10:**
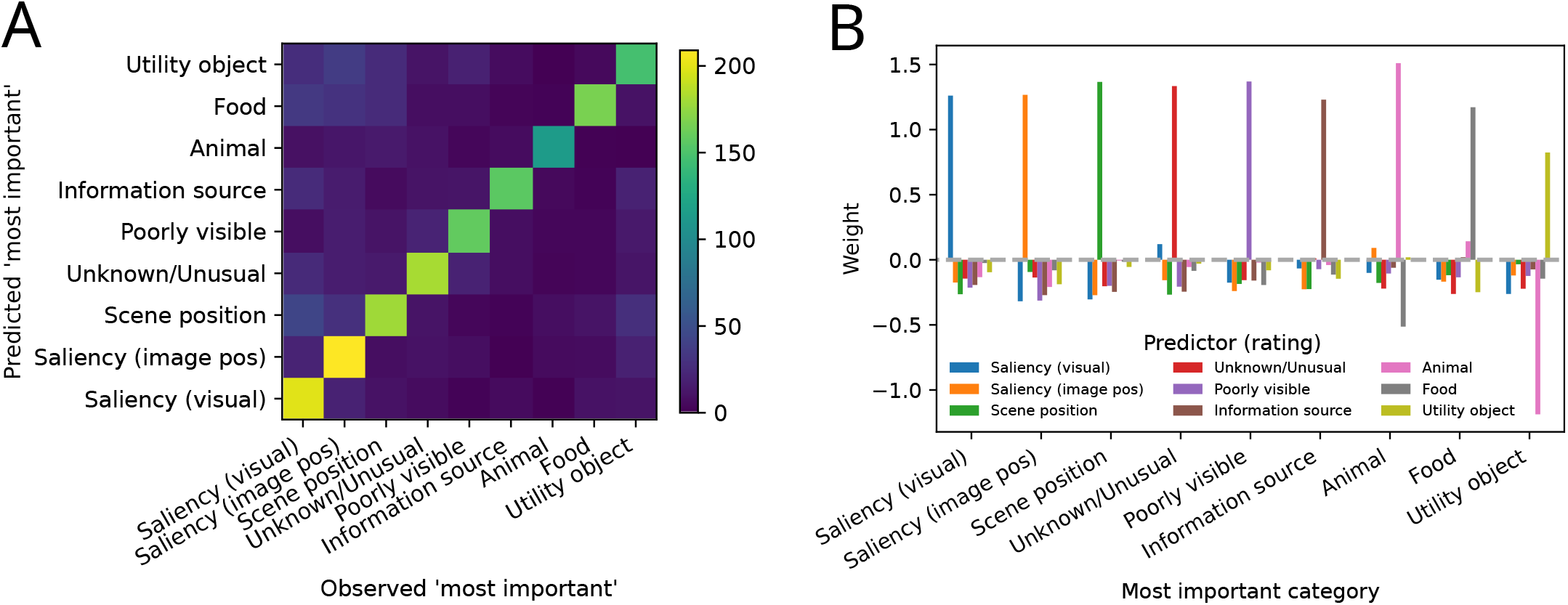
Results of a multi-class log-linear regression fit, used to predict ‘most important category’ choices from the category ratings. A: The confusion matrix, which plots the observed vs. the predicted ‘most important’ category. B: Regression weights of the best fit. For each of the most important categories, the nine bars show the weight of each category rating in the prediction. See text for details.

The model can also be used to predict the dominance matrix, and the fitted values are very similar to the observed dominance matrix (see Appendix A.1).

## 3 Relationship between direct and indirect methods

In the second part of the article, we clarify the relationship between the direct method proposed in the present work and existing indirect approaches for evaluating the relative importance of saliency and semantics. To support this discussion, Figure 11 presents a framework that outlines our assumptions about key components in gaze control and their potential interaction. Several elements of this framework relate to ideas that have been proposed in previous accounts of attention and eye-movement control in scenes (e.g., Henderson, 2020b; Navalpakkam & Itti, 2005; Sun et al., 2008; Wischnewski et al., 2010; Zelinsky & Bisley, 2015).

**Figure 11:**
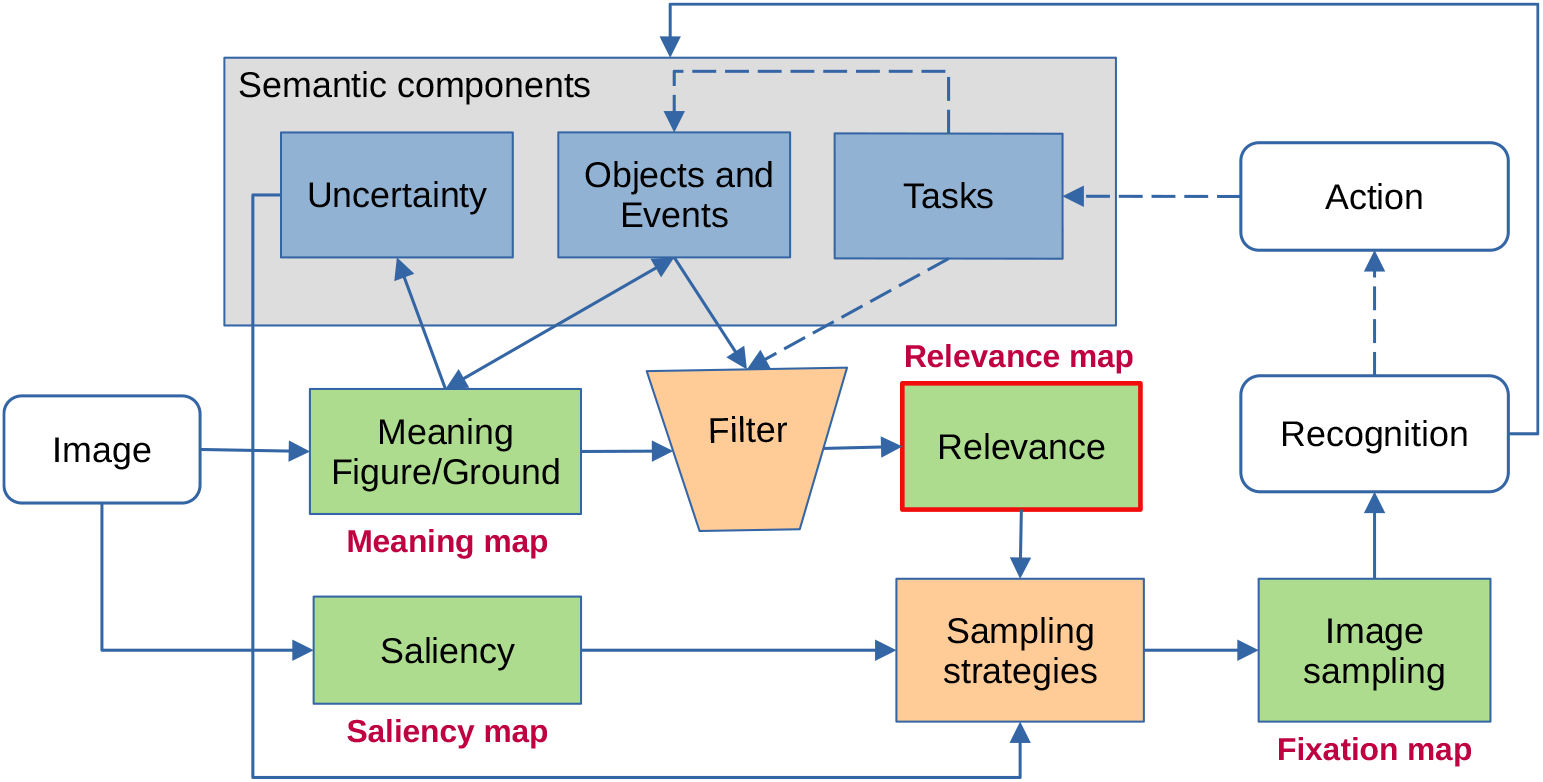
Scheme of key components contributing to gaze control and their proposed interplay. The scheme depicts a processing loop that typically works on a stream of input images. The processing starts with two early steps that result in a “saliency map,” where conspicuous locations are marked based on low-level image features, and an “object map,” in which potential object regions are outlined. The corresponding figure-ground segmentation draws on previous knowledge about objects and events. Image regions that cannot be classified produce local uncertainty. The “semantic map” is then weighted for importance, i.e., objects with high general relevance (e.g., faces or food) and objects that are useful for a given task, are emphasized according to their present relevance, whereas other object regions are attenuated. Information about regions where saliency, uncertainty, or relevance are high are then integrated and result in a sampling strategy, which plans the next fixation to a position that promises the highest information gain. Over time, the sampling strategy manifests itself in the fixation map. In a real world scenario, the information gathered at the fixation point would be used to recognize objects and to plan further action, which in turn could change input, scene knowledge, and task. In an experiment with free-viewing instruction, the image is static and actions and task play no role. This is indicated by dashed lines in the scheme.

In Figure 11, we highlight the representation termed “relevance map” as we consider it to be central for understanding the cognitive processes underlying object recognition and action planning in general and, more specifically, to explain the influence of semantics on gaze control. For a given visual input, this map indicates all objects that are relevant, either generally or for the task at hand. All other components may be regarded as just auxiliary processes that serve to build this map. Under normal viewing conditions, where only a small part of the scene is sharply visible and attended to, the relevance map cannot be generated in a single step, but must be constructed incrementally. We assume that this occurs via input sampling strategies that choose the direction and amplitudes of saccades in a way that minimizes the number of fixations required to extract the relevant scene content. This is supported by results from Murlidaran et al. (2026), which suggest that the “human free-viewing fixation patterns may emerge as a functional byproduct of optimizing scene comprehension under the biological constraints of foveated vision” (p. 1). These sampling strategies manifest themselves in the observable fixation pattern and are influenced by default strategies (e.g., a central bias), image saliency, uncertainties in the classification of the input and the current state of the relevance map.

In Figure 11, saliency is understood in its classic sense, that is, as a function *f* (*x, y, a*_1_, … *a*_*n*_) of the local center–surround differences in the responses of low-level feature analyzers *a*_*i*_ at location (*x, y*). The saliency map simply represents the joint distribution of these signals across the input (Itti et al., 1998; Itti, 2001). It is assumed that high saliency has a direct and automatic influence on the sampling strategy and increases fixation probability to the corresponding location.

Influences on the sampling strategy other than saliency are related to semantics in some form. They are more complex and draw on previous knowledge and the current task. If presented with a new input, a first step is to extract the gist of the scene (Greene & Oliva, 2009; Wolfe et al., 2011). An important aspect of this is to perform a figure/ground segmentation of the scene and to focus on the “figure” part (Russell et al., 2014). Typical background regions in indoor scenes are areas depicting walls, floors, curtains, or large furniture, whereas trees, vegetation, sky, lakes and sees play this role in outdoor scenes. The remaining “figure” regions are compared to known objects and filtered according to the a priori importance of certain object categories and— in action planing—according to their relevance for the task at hand (Li & Chen, 2021). Unknown objects or regions that cannot clearly be classified increase uncertainty that in turn may trigger attention and closer inspection (Chakraborty et al., 2022).

This distinction between saliency and semantics made here describes different processing stages, but it is likely not fundamental. From an evolutionary perspective, saliency is only advantageous if it is statistically linked to semantics; specifically, there must be a high probability that directing one’s gaze toward a salient location will reveal objects relevant to the observer’s survival or goals.

This schematic sketch may be used to highlight the specific aspects emphasized by different approaches to explain gaze control: One possibility realized in classic saliency models is to start from visual saliency and to try to predict observed fixations. These models focus primarily on sampling strategies that rapidly direct gaze to potentially interesting input locations (Masciocchi et al., 2009), whereas the complex influences of semantic information on fixations are not taken into account.

Meaning maps, as proposed by Henderson and Hayes (2017), can be regarded as the opposite approach, since they do not take saliency into account and instead can be interpreted as supporting a segmentation of the input into figure and ground. We assume here that the “meaning” ratings collected during the construction of meaning maps are better understood as reflecting the probability that a region belongs to a figure rather than the background. This appears more appropriate than an interpretation in terms of “meaning”, because meaning, understood as a driver for preferences or attentional priority, is at least partly subjective and context-dependent. Given this subjectivity and context-dependence, it appears difficult to distill ratings of meaning into a single, static map. Although this semantic segmentation captured by meaning maps ignores both differences in the subjective importance of object categories and any influences of task, it is a useful prerequisite for constructing the relevance map.

From this perspective, meaning maps appear to be related to the model of proto-object based saliency proposed by Russell et al. (2014). It should be noted that the implicit assumption that there exists a pre-attentive object representation is supported by a large body of evidence reviewed in Cavanagh et al. (2023). As in the saliency-map approach, observed fixations are not used in the construction of meaning maps. However, unlike saliency maps, meaning maps seem to have limited predictive value for fixations because their influence on the sampling strategy is not direct. According to our framework, it is instead mediated by a filtering operation that weights potential targets based on their current importance.

Machine-learning approaches, like DeepGaze IIE (Linardos et al., 2021), start from the fixations and, during training, try to learn the image properties that predict them. As fixation patterns are the manifestation of the sampling strategy, which according to our framework incorporate all influences from semantics and saliency, it is plausible that both saliency and all generalizable properties of objects and events, in particular those concerning their relative importance, can be learned and used in the predictions. In line with this assumption, such models often achieve more accurate predictions than those derived from saliency maps or meaning maps. From a cognitive science perspective, a key limitation of this machine-learning approach is that the information underlying these predictions remains largely implicit (Hayes & Henderson, 2021).

Like machine-learning approaches, the proposed direct approach relies on the observed fixation distribution, albeit only indirectly, for identifying fixation hotspots. However, the goal is not to predict fixations but rather to quantify the inputs to the sampling strategy and to identify how they are integrated. Thus, in a sense, the direct approach seeks to make explicit the information that machine-learning models (and human observers) use implicitly.

### Dissociation between fixations and important image regions

A key distinction made in the framework depicted in Figure 11 is between processes that indicate important regions in the input and a sampling strategy that generates fixations. This distinction is useful, because fixation locations generated by the sampling strategy do not cover subjectively important image regions uniformly and may also lie outside of these regions. As a consequence, one cannot always infer the relevance of image regions directly from fixation locations. This is important because several studies suggest that objects are the “unit” of attention (Z. Chen, 2012; Nuthmann & Henderson, 2010) and that objects predict gaze behavior better than local features (Borji & Tanner, 2016; Stoll et al., 2015).

We illustrate some reasons for this dissociation between fixations and relevant object regions with examples from our stimulus set, in which the “object of interest” is rather obvious (see Figure 12): (1) Approximate object recognition, although impaired in the periphery, is possible in a relatively large area around the fixation point. Thus, a single, well-chosen fixation point may suffice to fully recognize a large object. Typically, observers prefer a point near the center of the object or, if the object has different parts, a point near a part of special interest (for example, the head of an animal) [see panels B, F] (Nuthmann & Henderson, 2010; van der Linden et al., 2015; Yun et al., 2013). (2) For the same reason, a whole ensemble of objects may be recognizable with a single fixation, especially when they are distant in the scene and thus small and close to each other in the image. In this case, it may be optimal to fixate a point near the center of the object cluster even if this point itself contains no important information [see panel C] (Zelinsky et al., 1997). (3) If a relevant object is relatively small, fixations directed to this object may sometimes land outside its boundary [see panels A, C] (Pajak & Nuthmann, 2013). (4) Another reason why fixations sometimes lie outside important objects is an unequal distribution of information within an image region, because in this case a bias in fixation location toward higher information density may be advantageous. This is particularly relevant in eye-tracking experiments, where images are presented on a screen and relevant information drops to zero beyond the screen borders. A sensible strategy would be not to fixate an object near the border directly but to choose the largest distance to the border that still allows one to recognize the object [see panel E; the top of the region corresponds to the top border of the entire image]. (5) When objects move, it may be reasonable not to fixate the object center, but a position slightly shifted in the movement direction. This may even hold in static images, when the movement direction can be inferred [see panel D] (Açik et al., 2014). (6) Furthermore, fixations may also be largely unrelated to object relevance, for example when they are driven by saliency or uncertainty [see panel F]. In our experiment, such cases are indicated by regions in which visual saliency was named as the most important factor.

**Figure 12:**
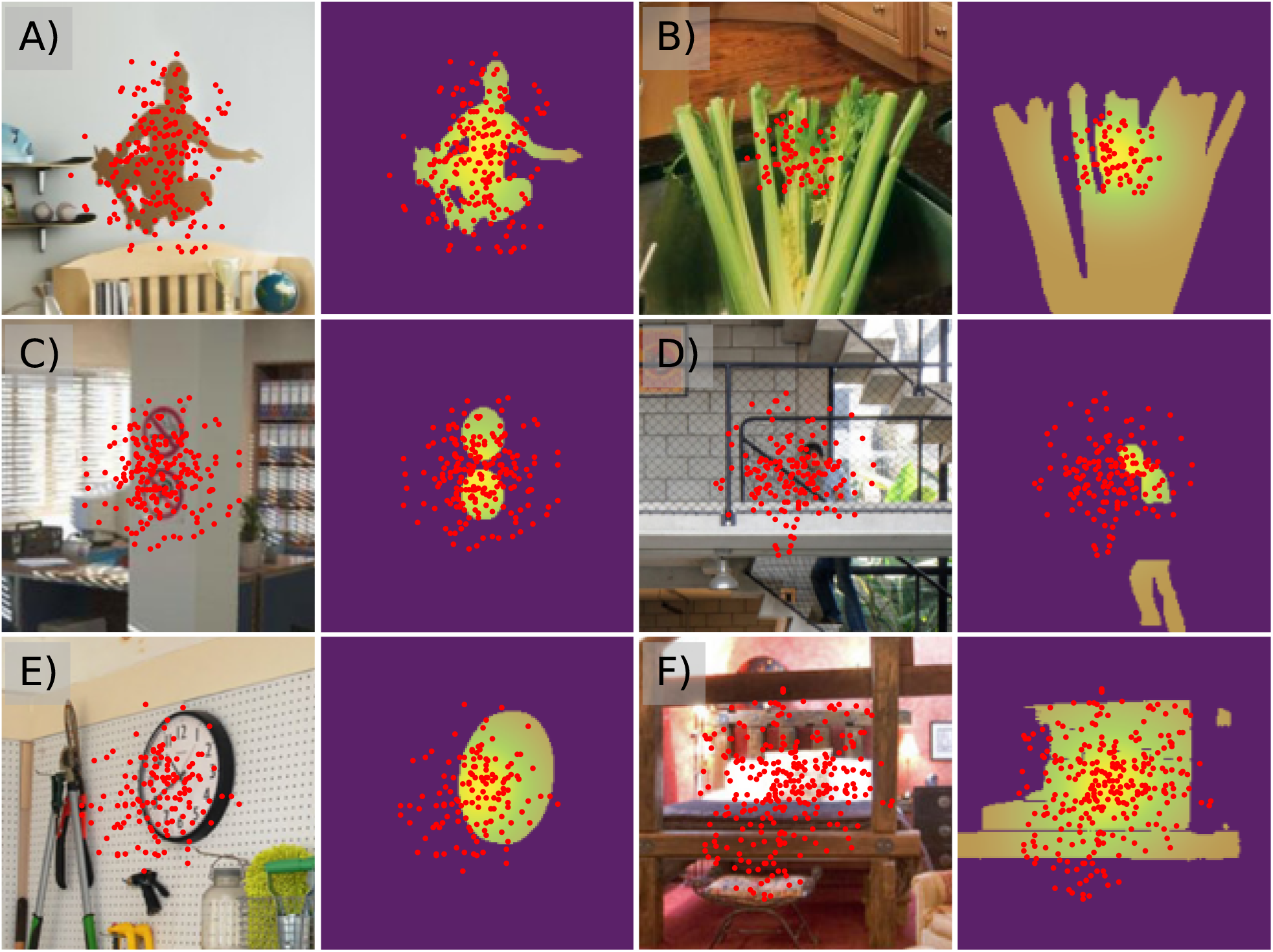
Examples from our data set, illustrating different causes for a partial dissociation between fixations and target objects: A) target region small, B) target region large, C) multiple targets, D) target motion, E) unequal information density, F) fixations not triggered by target objects (here ‘saliency’ was mentioned by our participants as the most probable cause). In each panel, the left image shows a subregion used in the experiment with overlaid fixation positions, and the right image shows a mask for the probable target objects overlaid with the estimated fixations distribution and the fixations itself. Note that, on average, each participant contributed only 2–3 fixations to the fixation distribution, i.e., the fixation pattern reflects similarities in gaze across participants.

### 3.1 Effects of the reason of fixation on prediction quality

In this section, we investigate whether and how information gathered with the direct method can be used to augment and strengthen conclusions drawn from other methods intended to predict and explain gaze behavior.

To this end, we consider meaning maps (Henderson & Hayes, 2017; Peacock et al., 2019), Adaptive Whitening Saliency (AWS) (Garcia-Diaz et al., 2012) as a representative for a classic saliency model, and the DeepGaze IIE model (Linardos et al., 2021) as a representative of approaches based on machine learning. We also include the empirical fixation map as a reference. In the case of AWS and DeepGaze IIE, the original code provided by the authors was used to create the maps. The meaning maps were generated as in Henderson and Hayes (2017), except that raters saw each image patch within its full-scene context, yielding contextualized meaning maps (Peacock et al., 2019). For the meaning maps and the AWS maps, the pixel-value histogram of each map was matched to the histogram of the continuous empirical fixation distribution for the same scene, ensuring that the overall value distributions of predicted and observed maps were comparable.

### Prediction of fixations

A common method to compare different saliency models is to test how well these models predict actual fixations (Kümmerer & Bethge, 2023). Recently, this approach was extended to pit meaning maps against saliency maps (Henderson & Hayes, 2017).

Several metrics have been proposed to quantify the degree of correspondence between predicted and observed fixation positions (Bylinskii et al., 2019). Among the measures discussed in that work, we selected three metrics that are representative of the diverse range of measures currently employed in the field: the Pearson correlation (CC), the Area under the ROC Curve (AUC), and Information Gain (IG). We report a normalized version of the standard AUC Judd score, calculated as AUC = (AUC_Judd-0.5)*2. Information gain quantifies the departure from spatial randomness and is calculated as the mean of the log likelihood ratio between the predicted fixation distribution and a uniform baseline distribution, where the average is taken over the observed fixation locations (Kümmerer et al., 2015).

The analyses were performed on 100 sub-images of size 200 × 200 pixels, centered at the 100 hotspots used in our experiment. For each sub-image, corresponding areas were cut from the prediction maps and only fixations inside the sub-images were kept.

Here, we explore the question to what extent the quality of the fixation prediction of a given map also depends on the reason for the fixation. To test this, we split the regions into subsets according to the semantic or saliency criterion that in our experiment was considered to be the most important reason for fixating a given region. However, we collapsed the nine categories into 5 by combining similar criteria: (1) ‘Object’ comprises the categories ‘utility’, ‘food’, ‘information source’, and ‘animal’, which all denote object classes of special importance for humans. (2) ‘Position’ comprises ‘image position’ and ‘scene position’, which both indicate that the main cause for the fixation is a prominent position either in the image or the scene. (3) ‘Saliency’ comprises only visual saliency, (4) ‘unknown’ stands for ‘unknown/unusual objects’, and (5) ‘visibility’ corresponds to the remaining category ‘poorly visible’.

The top row of Figure 13 summarizes the results. Since the chosen metrics assess the correspondence between the prediction maps and the ground truth (i.e., the observed fixation distribution), the empirical fixation map (Empfix) serves as an upper bound for the other predictions. This is most obvious for the Pearson correlation, where Empfix is correlated with itself. For the IG metric, which measures the information gain over a uniform distribution, the values for EmpFix vary slightly. This suggests that the concentration of the fixation distribution varies across the different fixation reasons, with the most concentrated distribution occurring in the ‘Saliency’ category.

**Figure 13:**
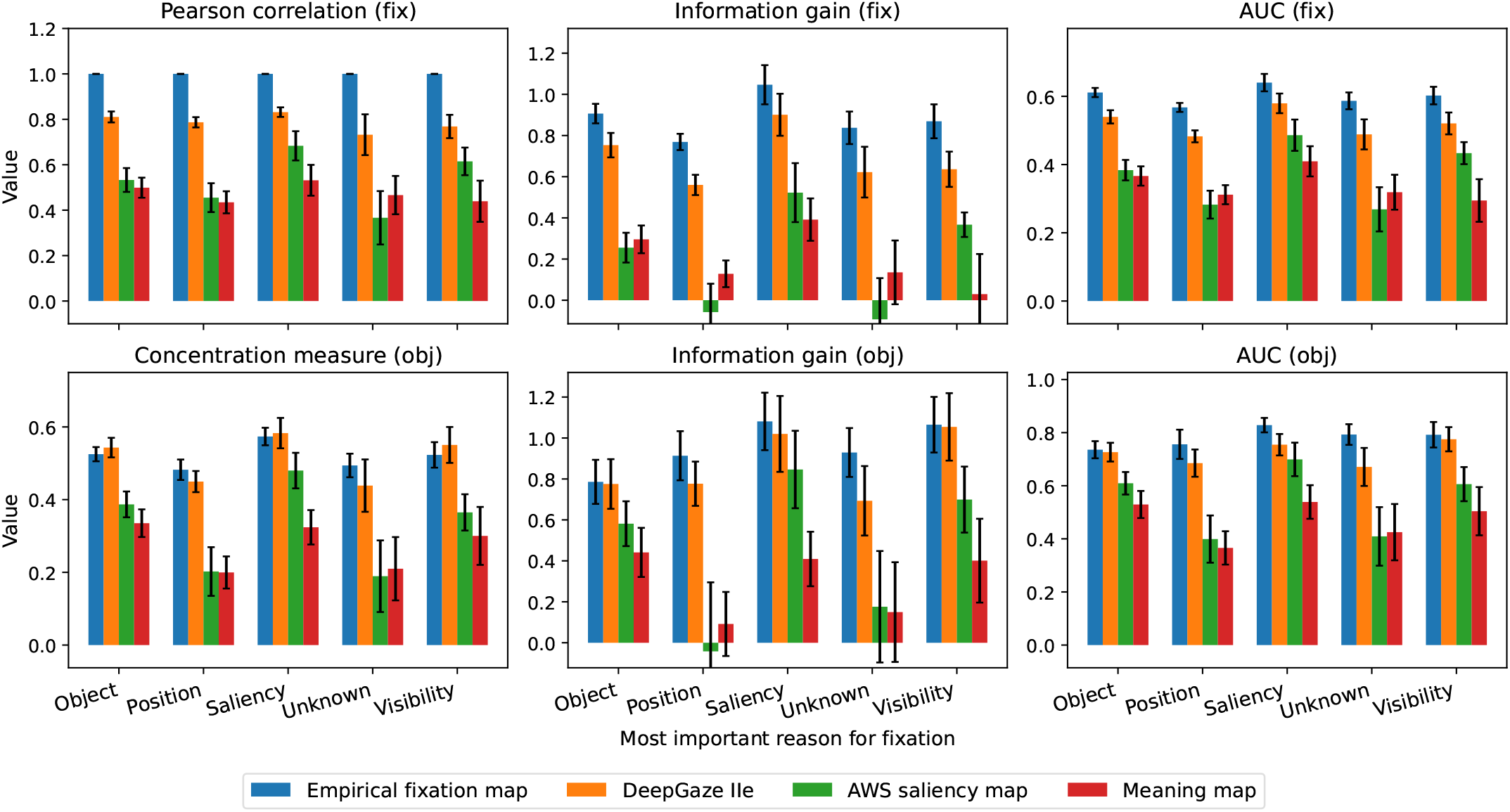
Quality of prediction of different map types for the 100 regions rated in our experiment, using a fixation-based criterion (top row) and an object-based criterion (bottom row). The map types are DeepGaze IIE maps, meaning maps, the AWS saliency maps, and, for reference, the observed fixation distribution (Empfix). The predictions are grouped according to the category which was rated as most important for the fixations in each region. Each row shows three different metrics. The errorbars enclose ± 1*SE*. See text for details.

Similar dependencies on fixation reason were obtained with all three metrics, but the differences seem slightly more pronounced with the IG measure. Overall, the DeepGaze IIE model made the best predictions, which are relatively close to the optimum and depend only slightly on the reasons for the fixation. In comparison, the predictions made by the AWS model are considerably less accurate and depend markedly on the reason for fixation. The latter is most obvious for the IG metric, where the information gains for ‘Position’ and ‘Unknown’ are close to zero or even negative. The overall low IG values for the AWS model indicate that it tends to make rather broad and unspecific predictions. In line with the theory underlying this model, the best predictions are made when saliency was rated as the most important reason for fixation. The performance of the meaning maps is on average similar to that of the AWS model, despite the differences underlying their construction. This is consistent with the findings of Pedziwiatr et al. (2021a). For meaning maps, the mean information gain over a uniform prediction is close to zero in the three categories ‘Position’, ‘Unknown’ and ‘Visibility’. This appears plausible, because neither of these factors is included in the construction of meaning maps.

### Predicting relevant object regions

Machine-learning approaches use observed fixations when training the model, whereas both meaning maps and classic saliency maps are constructed without any reference to fixation data. Instead, they rely solely on image information, with an implicit emphasis on the discernible objects in the scene (Elazary & Itti, 2008). For classic saliency models, there exists a direct theoretical link to fixation locations, namely the assumption that high saliency regions attract fixations. The situation is more complicated with meaning maps, because the fact that an image patch contains recognizable scene elements does not imply that it is also interesting and relevant for an observer. Evaluating the different models using metrics that measure how well fixations are predicted puts both AWS and meaning maps at a disadvantage. Exact fixation locations also do not seem to be the proper basis for relating our results to prediction maps, because although our approach starts with observed fixations to define regions of interest, the ratings in the experiment are mainly influenced by the properties of the *objects* within these regions.

To allow a fairer comparison, we constructed an estimate of the corresponding relevance map for each region image, containing only the scene elements to which the judgments in our experiment presumably refer (see Panel A in Figure 14 for an example). First, the GLIPv2 semantic segmentation model (Zhang et al., 2022) was applied to each region image. The prompts and segmentation thresholds for this procedure were manually optimized for each region to best approximate a “complete” segmentation of the image, where all objects were plausibly labeled. Next, we excluded regions with background labels (in our indoor scenes these are, for example, “wall,” “desk,” “curtain,” “shelf,” etc.), as these are almost never fixated during scene viewing without a specific task. Presumably, this intermediate representation captures information similar to what meaning maps encode. To further select the relevant target object from the set of possible targets, we used observed fixations inside the extended mask: only objects where either the number of fixations or the fixation density within their image region exceeded a certain threshold were retained. The union of the retained regions defines the uniform “relevance map” (see bottom left panel in Figure 14A). We also constructed a variant, in which the uniform relevance map is weighted with the fixation distribution (see bottom right panel in Figure 14A and Figure 12 for further examples).

**Figure 14:**
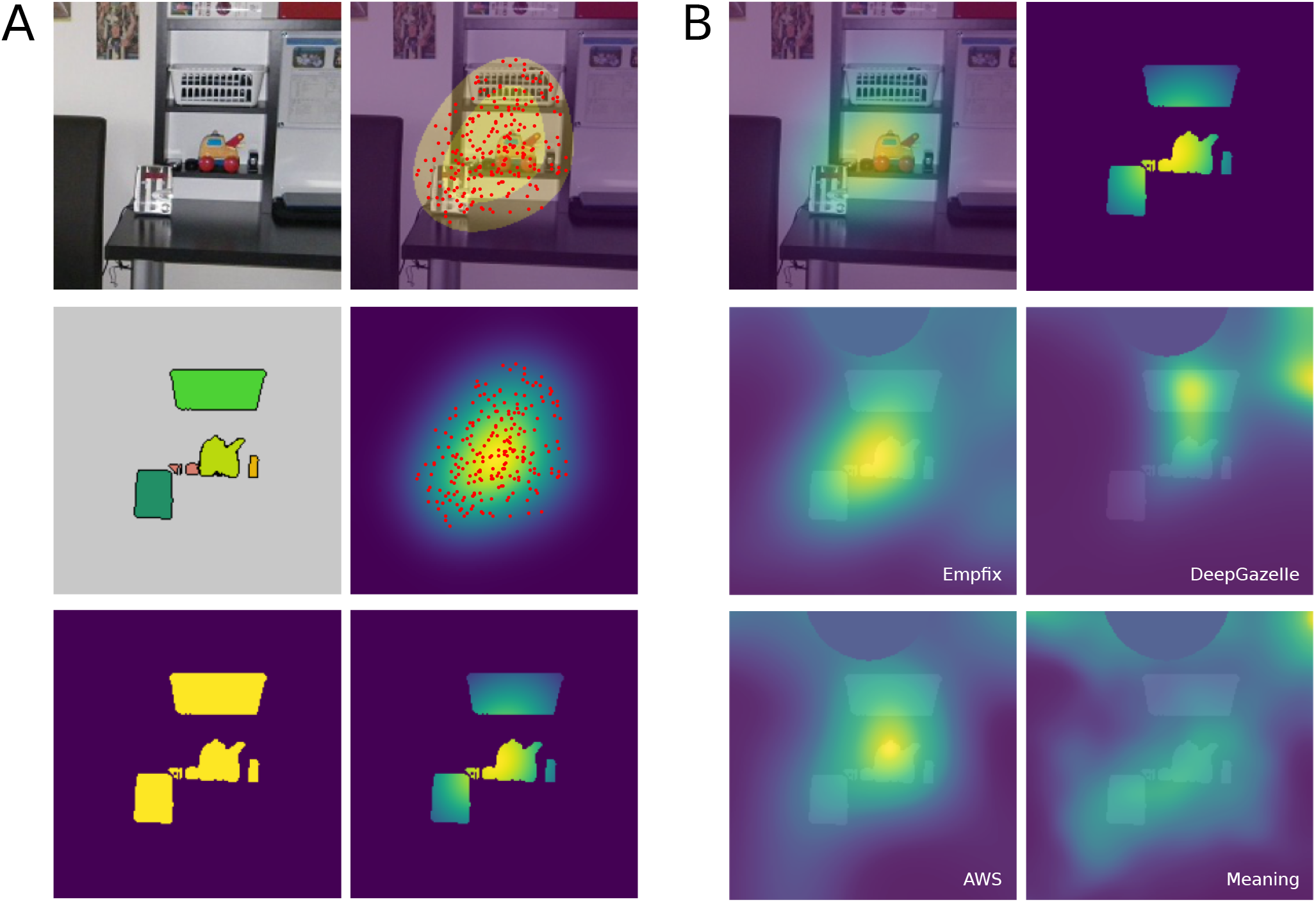
Panel A: Relevance map for one of our region images, estimated from combining semantic segmentation and fixation information. Top left: Region image. Top right: Mask and extended mask with overlaid fixations. Middle left: Non-background objects that were fixated. Middle right: Fixations in the extended mask and estimated fixation distribution. Bottom left: Uniform relevance map. Bottom right: Weighted relevance map, i.e., the uniform relevance map multiplied by the fixation distribution. Panel B: Visualization of the prediction maps for the region image shown in Panel A. Top left: Region image with overlaid fixation distribution. Top right: The weighted relevance map. Middle and bottom row: Uniform relevance map overlaid with a cutout of the four prediction maps. Note: In this case, the circular region at the top of each region image covers a neighboring region with high fixation density. The value of this region is set to the mean value in the rest of the map outside the uniform relevance map.

Panel B in Figure 14 shows the uniform relevance map overlaid with the four prediction maps for the corresponding region image. Again, three different metrics were used to evaluate how well the maps predict the relevance maps. In each case, the values of map *X* are interpreted as a probability distribution, i.e. Σ*P* (*X*) = 1 and *P* (*X*) ≥ 0. Instead of CC, we here use a Concentration Measure (CM), which relates the values *P* (*S*) inside the relevant object region to those outside *P* (*X*\*S*) : *CM* = (*P* (*S*) − *P* (*X*))/(*P* (*S*) + *P* (*X*). It has the same range [1, −1] as the Pearson correlation, where a value of 1 indicates perfect prediction, and a value of 0 equal probability mass in *S* and *X* \ *S*. The IG over a uniform distribution and the AUC measure are also applied in this case, simply by replacing the discrete fixation distribution with the uniform relevance map.

The bottom row of Figure 13 shows the corresponding values for each map type depending on the reason of fixation. Again, similar results are obtained with each of the three metrics. A general observation is that overall Empfix and DeepGaze IIE still make clearly better predictions than the two other models, but the advantage of Empfix over DeepGaze IIe is reduced or even vanishes. Furthermore, when compared with the results of the fixation-based criterion in the upper row, the differences between DeepGaze IIe and AWS are considerably reduced in the three categories ‘Object’, ‘Saliency’ and ‘Visibility’ and increased for the remaining categories ‘Position’ and ‘Unknown’. A similar result is observed for meaning maps, but to a lesser extent. This is plausible for ‘Saliency’, which plays no special role in meaning maps.

These results for predicting relevant objects indicate that one reason why machine-learning approaches outperform classic saliency models and meaning maps in predicting fixations is their ability to learn not only object-related information but also other factors influencing the sampling strategy. Additionally, the findings suggest that the quality of object prediction varies depending on the reason for fixation. When saliency was the primary driver of fixation, the performance of AWS was comparable to that of DeepGaze IIE. Somewhat surprisingly, meaning maps showed limited improvement when relevant objects are considered instead of fixations. One possible explanation for this could be the relatively low spatial resolution of meaning maps, which is a consequence of the fact that the patches used in their construction are of fixed size and relatively large (Pedziwiatr et al., 2021b).

## 4 General Discussion

The present work was motivated by potential limitations of studies that seek to determine whether saliency or semantics plays a greater role in gaze control by comparing the predictive power of saliency maps versus meaning maps for observed fixations (e.g., Henderson & Hayes, 2017; Peacock et al., 2019).

The logic underlying this comparison follows an established practice: first, construct a spatial map representing a putatively relevant factor (e.g., contrast, saliency, surprise, or meaning) based on specific theoretical principles; second, evaluate the validity of those theoretical principles by assessing how well the resulting map predicts observed fixations. Within this approach, after appropriate normalization, the maps are typically interpreted as predicted fixation probability densities.

In the case of saliency maps, where saliency is assumed to directly attract attention, high map values plausibly correspond to high fixation probabilities. For meaning maps, however, the link between the operational definition of meaning and fixation probability is much less direct. For example, highly recognizable objects are often of little interest in themselves and may attract attention only when they appear in unexpected contexts or locations. Conversely, among objects of comparable recognizability, some such as faces are inherently more attention-grabbing.

### Experiment: Identifying reasons for fixations

A further limitation of this approach is its implicit assumption that either semantics or saliency exerts a dominant influence on gaze control globally. However, visual inspection of preferentially fixated image regions suggests that fixations arise from heterogeneous causes, with the primary influence varying locally within a scene and across scenes.

To test this hypothesis, we employed a direct approach to identify the reasons for fixations. We first used existing fixation data for a set of scenes to locate hotspots within each scene. In a subsequent experiment, participants saw each scene with a single outlined hotspot derived from these data. The participant’s task was to first rate the extent to which two low-level saliency categories and seven semantic categories described the hotspot’s content. In a second step, they selected the one category they considered most responsible for attracting fixation. In total, 100 hotspots (two from each of 50 scenes) were rated. Although the raters were untrained, the ratings were highly reliable. These results indicate that it is feasible to both quantify the presence of potential fixation determinants and identify the most probable one. Moreover, for each category, the probability of being selected as the primary reason increased monotonically with its rating, consistent with the interpretation that these categories represent genuine determinants of fixation.

In the analysis, we considered all categories with a rating of at least 2 (on a scale from 0 to 5) as relevant, or briefly, as “active.” A first observation was that, in most cases, several categories were active, indicating that multiple factors may have contributed to a single fixation. Because each participant had to select a single category as the most important one, the contribution of multiple categories can be inferred only indirectly from the distribution of choices across observers. Approximately three categories are needed to account for more than 80% of observer selections of the most important category (Figure 3C).

Both saliency categories, “visual saliency” and “preferred image position”, were active in over 50% of cases, confirming that fixated regions are often salient. However, these types of saliency do not appear to be necessary conditions for fixation, as both categories were inactive in more than 15% of hotspots.

Regarding semantics, we distinguish between object-based and abstract categories. Three of the four object-related categories, “information source,” “animal,” and “food,” correspond to object classes well-established as frequently fixated (Li & Chen, 2021). These three are mutually exclusive and, accordingly, were found to correlate negatively with each other. The remaining object-related category, “utility object,” is broader in scope and encompasses most man-made objects with which people interact, such as tools, dishes, or electronic devices. Object-related categories encompass those with highest relative importance, with “animal” and “food” being especially important. Consistent with the broad scope of the “utility object” category, it was the most frequently active among the semantic categories. At the same time, however, it was assigned the lowest importance across all categories. This partly reflects the fact that the “utility object” category overlaps with other, more specific categories (e.g., an alarm clock is both a utility object and an information source) or co-occurs with them in the same location (e.g., food on a plate). In these cases, the more specific category is usually preferred. Nevertheless, the discrepancy between high activation frequency and low assigned importance also supports the view that assigning high meaning to an object when creating a meaning map does not necessarily indicate a high probability of fixation.

The three abstract categories were “prominent position in the scene,” “unknown or unusual object,” and “poorly visible.” Among these, the “unknown or unusual object” category was the most important, nearly as important as the “food” and “animal” categories. This suggests that observers find novel or unidentifiable objects especially interesting, highlighting the role of prior experience in predicting the allocation of attention and gaze. While one might intuitively expect viewers to prefer personally relevant objects, our findings indicate that this preference may be superseded by the presence of novel stimuli. This confirms the result of Chakraborty et al. (2022), which used an operational definition of recognition uncertainty that focuses on the number of object categories competing for an object.

The “poorly visible” category was rated as less important than the “unknown” category but still more important than either saliency category. The strong positive correlation between “poorly visible” and “unknown or unusual object” suggests that some items could not be recognized due to suboptimal viewing conditions. Nevertheless, the two categories behaved differently: for instance, “poor visibility” correlated negatively with “visual saliency,” whereas “unknown or unusual object” correlated positively, indicating that these categories capture distinct aspects of fixation behavior.

The third abstract category, “prominent position in the scene,” was intended to capture cases in which an object attracts attention due to its location within the scene rather than its intrinsic properties—for example, an object placed within a spotlight or on a visually prominent surface (e.g., a central table). However, inter-rater reliability for this category was low, indicating that participants interpreted the definition inconsistently and suggesting that the instructions were ambiguous. A further complication is that this category represents a hybrid between saliency-based and semantic-based criteria. It is semantic in that it requires scene-level understanding and segmentation (e.g., identifying what constitutes a “conspicuous” or “privileged” location). Yet, like saliency-driven attention, it directs gaze to an object independently of the object’s identity or meaning, relying instead on contextual or spatial cues. This dual nature may have contributed to the ambiguity participants experienced when applying the category.

### Saliency vs. semantics

The central question motivating this study was whether visual saliency or semantic information more strongly guides gaze control. Our results reveal that the determinants of fixation are heterogeneous across a scene; consequently, any assessment of saliency versus semantics can only be made at the level of individual fixation hotspots. When results are aggregated across hotspots, the outcome inevitably reflects the particular composition of the scene: in abstract scenes lacking strong semantic cues, saliency should dominate, whereas in images populated with objects that are clearly recognizable but have a low contrast to their surround, semantic information is likely to prevail. Thus, the method employed by Henderson and colleagues (Henderson & Hayes, 2017; Peacock et al., 2019), which compares whole-scene saliency and semantic maps to infer the dominant cue, is fundamentally limited because it conflates heterogeneous local contributions into a single global metric (see also Murlidaran & Eckstein, 2025).

Despite these reservations about aggregation, our method of local measurements allows us to pose the saliency-versus-semantics question in a more nuanced way, by analyzing the relationship between category ratings and reported importance *within* each hotspot. We grouped the seven semantic and two saliency categories into corresponding super-categories and used them to analyze both the highest relevance ratings and the selection of the most important category. In trials where participants’ highest rating and their choice of the most important category were consistent (i.e., both belonged to the same super-category), semantics dominated (78.1% semantic vs. 21.9% saliency). The more informative evidence, however, comes from the inconsistent trials, where the highest-rated category and the most important category were from different super-categories. Here, participants selected a semantic category as most important in 74.1% of cases, despite the fact that a saliency category had received the highest relevance rating, whereas the converse was only true in 25.9% of cases. This preference for semantics in ambiguous situations indicates that semantic content is a more fundamental determinant of perceived importance than low-level saliency. Furthermore, the ubiquity of semantic information is underscored by our finding that in virtually all trials at least one semantic category was rated highly, whereas both saliency categories were inactive in over 15% of cases.

Taken together, these results demonstrate that the determinants of gaze control are intrinsically local and heterogeneous: the primary driver of a fixation can be a low-level saliency feature at one hotspot and a specific semantic attribute at another. This view explains why a global assessment of the importance of saliency and semantics can lead to inconsistent results depending on the stimulus material. Furthermore, our analysis reveals that the influence of semantic information is not monolithic, but rather composed of various distinct factors. While object-based semantics is an important influence, more abstract visual categories also play an important role. In particular, the “unknown or unusual” category proved to be relevant, underscoring the importance of individual experiences and prior knowledge in gaze control. Furthermore, the boundary between saliency and semantics is sometimes blurred—for example, in our category “prominent position in the scene,” which represents a hybrid case.

### Relationship between direct and indirect approaches

In the second part of the paper, we sought to relate the data obtained using our method to the types of information represented in existing approaches to explain fixation patterns during free viewing and scene-viewing tasks, where the task establishes general viewing goals (e.g., memorization). Specifically, we considered a classic low-level saliency model (AWS), a learning-based saliency model (DeepGaze IIE), and meaning maps. Although all three approaches lead to spatial maps that aim to predict fixations, they differ markedly in their theoretical foundation, their predictive scope, and the data on which they are based.

Figure 11 summarizes our view of the information flow underlying gaze control and situates the different map types within this framework. In this scheme, we made a key distinction between a) processes that signal potentially relevant locations in the visual input and b) a sampling strategy that converts this information into a sequence of fixations.We consider this distinction as important, because a detailed inspection of actual fixation patterns suggests that fixations are not always directed toward the most informative or salient positions in the scene. Rather, the sampling strategy appears to optimizes saccade vectors in order to minimizes the number of fixations required to capture relevant information in the scene(Murlidaran & Eckstein, 2025; Murlidaran et al., 2026). Some possible optimization criteria used in allocating fixations are illustrated in Figure 12. The sampling strategy can thus be conceived as a process that integrates information from previous fixations with cues to potentially informative next locations, selecting subsequent fixations to maximize information gain.

Within this framework, classic low-level saliency can be understood as a relatively simple mechanism that identifies visually prominent locations and passes them directly to the sampling strategy. In the context of our experiment, this corresponds closely to the category “visual saliency,” whereas the second saliency category “preferred position in the image,” represents a spatial bias intrinsic to the sampling strategy itself. All other sources of information are assumed to be related to semantics in some form.

We consider meaning maps as primarily reflecting a kind of figure-ground segmentation process, in which uninformative background regions are discounted. To perform such a segmentation, at least some degree of semantic understanding is required. The extent to which the remaining “figure” regions, typically corresponding to objects or “proto-objects” (Y. Chen & Zelinsky, 2019; Nuthmann & Henderson, 2010; Russell et al., 2014), are considered relevant at a given moment depends on an additional filtering process that reflects task demands and/or individual preferences. The object-based categories used in our experiment were designed to capture such intrinsic preferences. This filtered semantic information then serves as input to the sampling strategy.

The abstract semantic categories “unknown/unusual object” and “poor visibility” used in our study are functionally similar to saliency in that “figure” regions that cannot be readily recognized are assumed to provide direct input to the sampling strategy.

The data obtained with our direct method are assumed to characterize different types of inputs to the sampling strategy. Similar to classic saliency and meaning maps, this information is derived directly from the image, without reference to the actual fixations, which in our framework constitute the *output* of the sampling strategy. In contrast, learning-based saliency models start from the observed fixation data and, during training, attempt to identify the image features that best predict them. As a result, the trained model implicitly encodes not only the inputs to the sampling strategy but also aspects of the strategy itself. Whether this constitutes an advantage depends on the intended purpose. For predicting fixation locations, such implicit encoding should improve performance and this has been consistently observed (Pedziwiatr et al., 2021a). However, if the goal is to identify the attended objects or the underlying cognitive determinants of gaze, the predicted fixation positions may be misleading, as they partly reflect learned properties of the sampling strategy rather than purely stimulus-driven inputs.

In order to estimate how much of the superiority of learning based models over meaning maps and saliency maps in predicting fixations is due to knowledge of the sampling strategy, we compared a fixation-based and an object-based criterion of prediction quality. The latter requires knowing which objects the observers were interested in. In our study, this was relatively straightforward, as we focused on isolated hot spots in the fixation maps. With the usual fixation-based criterion, map values at fixation positions are compared with those at non-fixated areas. Analogously, with the object-based criterion, map values inside the objects are compared with values outside.

When using the object-based criterion, the superiority of the learning-based model often decreases somewhat, especially in hotspots where the reason for fixation corresponds to the information contained in the other two map types. However, the overall pattern suggests that this is not the only, and probably not the most important, factor underlying the performance of the learning-based model. In this context, another observation is that, with the object-based criterion, the predictions of the learning-based model are even closer to the predictions with the observed fixations than with the fixation-based criterion. This suggests that the deviations from the gold standard, i.e. observed fixations, found with the fixation-based criterion are partly due to the incomplete inclusion of actual sampling strategies in DeepGaze IIE.

### Reasons for fixation and prediction quality

Classic saliency maps are designed to capture visually conspicuous regions, whereas meaning maps focus on semantic information. In our framework, this difference is reflected by the specific location of each map type in the overall information flow. Learning-based saliency models, in contrast, are not per se limited to a particular type of information and may even partly incorporate aspects of the sampling strategy.

The results of our direct method allowed the classification of the 100 hotspots according to the most important reason for fixation. This made it possible to test whether the fixation prediction by each map is particularly strong when the most important cause of fixation in a hotspot corresponds to the information captured by the map. Specifically, we expected to find that classic saliency models excel if saliency was rated as the most important reason for fixation, whereas meaning maps should perform better if semantic criteria were rated as more relevant. Learning-based models should always show superior performance, especially if the dominant fixation cause belongs to a category not captured by the other maps.

It should be noted that with respect to a single map type this is not a particular strong test because, as our results show, there are usually several different potential causes of fixation. That is, even when the contribution of the most important factor is missing in the map, attention may nevertheless be guided by another slightly less relevant factor that *is* present. Furthermore, only a few scenes belong to each category. Nevertheless, the method seems useful to compare the performance of different map types.

Our results confirm that the performance of AWS, a classic saliency model, is particularly strong when the most important reason for fixation was ascribed to saliency, and particularly weak when positional information or an unknown object was selected as the main determinant of fixation. The performance of the AWS model was also relatively good for object-based semantic categories and, surprisingly, for the category “poor visibility”. Our results suggest that many objects that carry special meaning, for example food or faces, are also visually salient (see left panel in Figure 6; see also Nuthmann et al., 2020). Why AWS also performed well for regions with low visibility is less obvious, because in our experiment low visibility correlated negatively with saliency. A possible explanation is that AWS, which relies only on low-level information, still classifies some of these regions as salient, for example, dark regions due to shadows that stand out from the brighter surrounding, or text that is not legible but still exhibits large contrasts.

The performance of the meaning maps was often very similar to that of the AWS maps, as shown by Pedziwiatr et al. (2021a), but in some specific cases clearly worse. In line with the underlying design principle, when saliency was the main factor, prediction performance was lower than that of the AWS map. Hotspot where “poor visibility” was the main factor were also less well predicted by meaning maps than by AWS, which is plausible, because such regions should get low meaning ratings in the generation of the map. In our analysis, which only considered image regions that were frequently fixated, AWS overall performed slightly better than meaning maps.

As expected, the absolute performance level of the deep learning-based model DeepGaze IIE was much higher than that of the other two models, but in addition the performance also varied much less across different causes of fixation. Even in cases where the categories “position” or “unknown” were selected as the most import factors for fixation, the model performed well. This shows that the poor performance of both AWS and meaning maps in these cases can be attributed to a failure to capture important factors driving fixation behavior.

These results again demonstrate that the comparisons of the relative performance of different models depends on the actual causes of the fixation and thus on the specific scenes used in the benchmark. This is in line with the finding that the performance of saliency models varies substantially across scenes (Nuthmann et al., 2017).

### Limitations of the direct approach

Although the proposed direct method produced reliable and consistent judgments concerning the causes of fixation, there are theoretical and practical limitations.

A first critical point concerns the assumption that, for a given hotspot in the fixation distribution, participants can infer the underlying reason for fixation. While this assumption appears generally plausible and is supported by the high inter-rater reliability observed in our study, such judgments may nonetheless underestimate the role of visual saliency. This limitation arises because, during the evaluation, raters always fixate directly on the corresponding image region. In the actual viewing situation, however, a region may have been located in the periphery of the visual field at some position in the scanpath, making the image content less recognizable. Under such conditions, saliency may play a more prominent role than suggested by the rating and could even be the primary factor guiding attention.

Furthermore, the method requires a predefined set of potential reasons for fixation. This set must be sufficiently comprehensive to cover most cases that actually occur within the stimulus material, yet limited enough to remain practically manageable in terms of time and resources. In our study, we sought to mitigate this issue by restricting the range of scene types included. Another challenge lies in formulating concise category descriptions that are sufficiently clear and unambiguous to allow for reliable and consistent ratings across participants.

In our experimental paradigm, participants were asked to select a single category that they considered most important. Although this approach has certain advantages—most notably, it reduces participant effort— it also entails a loss of information regarding the relative importance of the categories. We sought to partially recover this information by examining the choice distribution across participants, but a more direct assessment could be useful.

A more practical concern, rather than a limitation of the method itself, is that the rating process is time-consuming, which limits the number of hotspots that can be evaluated within a single experiment. In such cases, recruiting participants through crowdsourcing platforms may be a practical option.

### Summary and conclusions

Our study contributes to the ongoing debate on whether saliency or semantics plays a larger role in gaze control.

We proposed and evaluated a method to answer this question based on direct judgments of the reasons for fixations at positions in the scene that are known to be frequently fixated.

Such judgments were found to be reliable and consistent across observers. They show that multiple factors contribute to gaze control and that the relative importance of each factor, in particular the relative contribution of saliency and semantics, varies considerably across the scene. This means that approaches that try to derive a global measure for the relative importance of both factors are limited in scope.

Still, we present some empirical findings and theoretical arguments supporting the conclusion drawn from previous findings that semantic information plays a dominant role in gaze control.

We present a conceptual framework of the information flow during gaze control that distinguishes between different processes indicating relevant regions in the image and a sampling strategy that integrates and weights this information and generates sequences of fixations. This framework helps to clarify how classic low-level saliency maps, learning-based saliency maps, and meaning maps, as well as our method, relate to each other and which specific information they capture.

We further provide empirical evidence that the performance of the three types of maps in predicting fixations depends differently on the cause of fixation. This indicates a specific reason why the relative performance of different models in benchmarks may depend on the stimulus material.

Further evidence suggests that the superior performance of learning-based models, which use empirical fixation information during training, is partly due to their incorporation of properties of the sampling strategy.

## A Appendix

### A.1 Simulation of the dominance matrix using the log-linear model

To compute the dominance between two categories *c*_*i*_ and *c* _*j*_, we initialize the mean rating vector *m*_0_ of length 9 to zero, and set the values at positions *i* and *j* to *m*_*i*_ and *m* _*j*_, respectively, where *m* is the mean rating for the corresponding category. The model is then used to predict the probabilities *p*_*i*_ and *p* _*j*_ of both categories (using procedure model.predict_proba(*m*_0_) of the fitted model). The dominance *d*_*ij*_ of *c*_*i*_ over *c* _*j*_ is then given by *d*_*ij*_ = *p*_*i*_/ (*p*_*i*_ +*p* _*j*_), and *d* _*ji*_ = *p* _*j*_/ (*p*_*i*_+ *p* _*j*_) = 1 − *d*_*ij*_. The result depends not only on the model, but also on the choice of *m*. One possibility is to use the observed mean ratings across all conditions. This works reasonably well, if rating < 1 are excluded. Otherwise, the relation between means for different categories depends also on the relative frequency, with which the categories are observed. The corresponding result is shown in the left panel of Figure 15. Another possibility is to search for the mean vector that minimizes the squared difference to the observed dominance matrix. The corresponding result obtained using global optimization (computed with scipy.optimize.differential_evolution) is shown in the right panel of Figure 15. Both choices lead to a similar dominance pattern, and the dominance matrix with optimized mean vector is almost identical to the observed dominance matrix shown in Figure 16.

**Figure 15:**
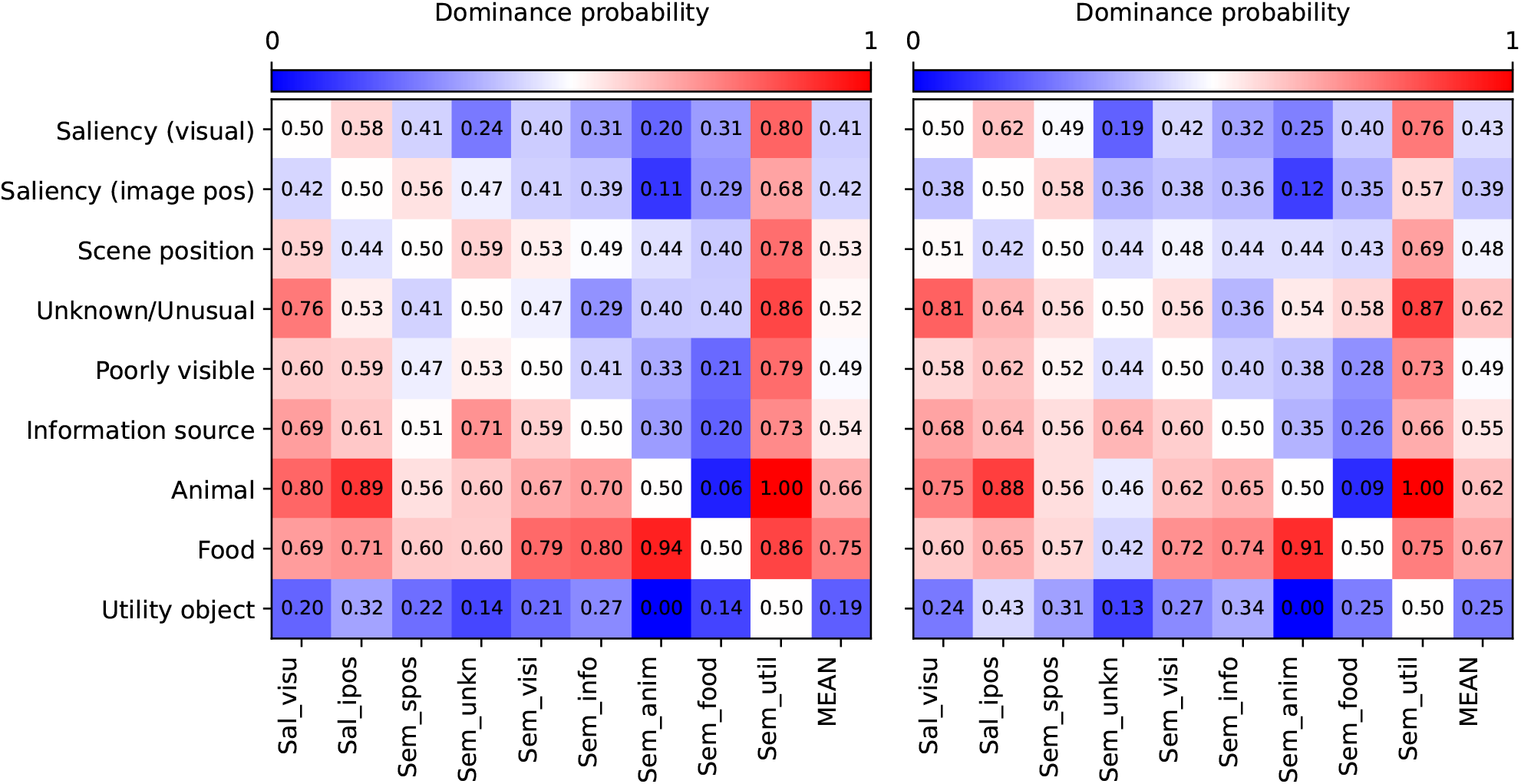
Dominance probabilities estimated from the loglinear model, using the observed mean of ratings > 1 (left), or mean ratings that minimize the squared distance of the observed dominance matrix.

**Figure 16:**
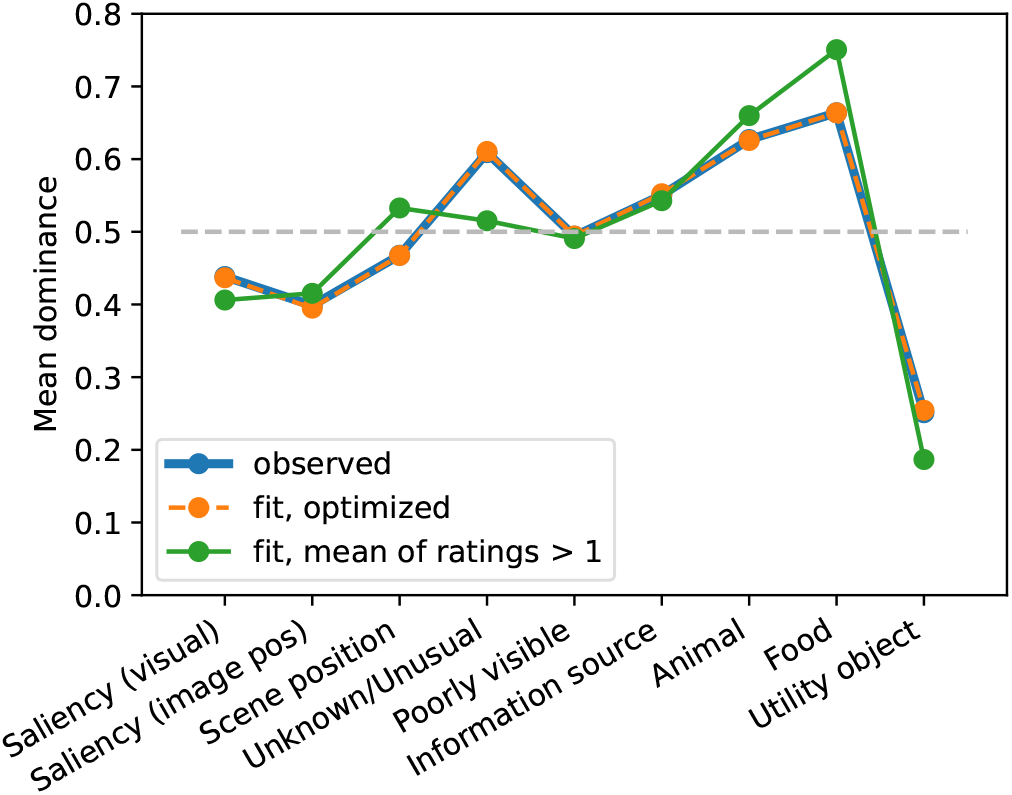
Observed mean dominance vs. fitted mean dominance. The observed mean dominance is very similar to the mean dominance resulting from the regression model.

